# Hybrid Semantic–Lexical Retrieval for Source-to-CDE Mapping in the NCI caDSR

**DOI:** 10.64898/2026.10.14.758967

**Authors:** James H. Tanis, Denise B. Warzel, Rakesh P. Khanna, Qingrong Chen, Chunhua Yan, Ravi Velleleth, Matthew T. Nash, Deepali Manathara, Brad B. Biggers, Joyce B. Soares, Daoud Meerzaman

## Abstract

Accurate mapping of local source data elements to standardized common data elements (CDEs) is critical for biomedical data integration but is challenged by short, abbreviated, and heterogeneous source descriptions. We constructed a benchmark of approximately 69,000 expert-linked source-to-CDE mappings from the National Cancer Institute’s caDSR and evaluated multiple bi-encoder models with two-stage fine-tuning. Here we developed a hybrid retrieval system combining semantic bi-encoder retrieval, CDE Match-inspired keyword retrieval, cross-encoder scoring, and supervised reranking. This system achieved Recall@5 ranging from 0.802 to 0.972 across internal and five distribution-shifted holdouts, outperforming the other evaluated methods on five of six datasets and approaching ceiling performance on the sixth dataset (GDC). These results demonstrate the effectiveness and robustness of integrating semantic and lexical evidence for source-to-CDE mapping, supporting improved interoperability and reuse of biomedical metadata across diverse clinical and research contexts.

## 1 Introduction

Organizations such as hospitals, research networks, and clinical studies often collect data using local naming conventions rather than a shared metadata standard. As a result, the same concept may appear under different variable names, labels, or question text across datasets. Mapping these local source data elements (SDEs) to standardized common data elements (CDEs) is therefore an important step for data harmonization, interoperability, and cross-study reuse [1, 2]. SDEs are local metadata fields—variable names, form labels and question text—used in clinical studies, registries and electronic data-collection systems, including EHR-derived workflows; the inputs to this task are therefore metadata labels and question text, not patient-level records. NCI CDEs are the standardized definitions used to keep those fields consistent across trials and studies [2, 3].

In practice, this mapping task is difficult. Human experts ultimately determine the correct match, but manual review is time-consuming and relies heavily on a priori knowledge of NCI CDEs, particularly given the large candidate search space. The production candidate catalog against which queries are ranked contains roughly 63,000 non-retired CDE records. Benchmark construction and production retrieval use different caDSR snapshots, which we describe in Section 2.1. Curators therefore depend on ranking systems to surface likely matches from source-query text derived from SDE metadata. However, source-side queries are often short, abbreviated, heterogeneous in style, and semantically underspecified. As a result, methods based primarily on lexical overlap may miss correct mappings when wording differs between the source element and the target CDE [4, 5].

We constructed a benchmark of approximately 69,000 expert-linked source-to-CDE mappings from the National Cancer Institute’s Cancer Data Standards Repository and Registry (caDSR) CDEs and used it to evaluate bi-encoders, text representations, and two-stage fine-tuning against a production catalog of approximately 63,000 CDE records. For controlled experiments, we used a local Python implementation developed from the service’s supplied source code that reproduces its core rule cascade and scoring logic against a frozen catalog, although minor implementation and data differences can affect individual rankings.

Earlier metadata-mapping systems relied mainly on lexical and terminology-based methods. Mougin et al. mapped source metadata to reference resources using lexical matching and field values [4], while Pathak et al. combined lexical and ontology-based methods across eMERGE institutions [5]. More recently, Fan et al. combined exact, lexical, and embedding-based semantic matching in a semi-automated pipeline for multi-site EHR flowsheet alignment and concept mapping [6], while CDEMapper combined Elasticsearch, GPT-based embeddings, and reranking for NIH CDE recommendation [7]. In parallel, bi-encoders have enabled semantic retrieval when relevant items share few exact terms with the query [8, 9, 10], including biomedical models such as PubMedBERT, SapBERT, and BioSimCSE [11, 12, 13]. Hybrid sparse–dense retrieval can improve over either representation alone [14], and multi-stage systems can apply cross-encoders to rerank merged candidate sets [15, 16].

These developments make bi-encoders a natural candidate for source-to-CDE retrieval, where the correct CDE may express the same concept as the query using substantially different wording. Task-specific and domain-adaptive training have improved bi-encoder representations in other retrieval settings [10, 17, 18, 19], but, to our knowledge, have not been evaluated for source-to-caDSR CDE retrieval. We therefore test whether fine-tuning on expert-linked source-to-CDE pairs improves retrieval.

Dense semantic retrieval alone, however, captures only part of the evidence useful for this task. Bi-encoders capture overall semantic similarity, whereas the correct mapping can depend on fine-grained lexical evidence. An abbreviation, qualifier, identifier fragment, or rare technical term may distinguish otherwise similar CDEs. Lexical methods preserve these cues [14, 20] but can miss correct mappings when the query and target express the same concept using different words [4, 5].

We therefore combine semantic and keyword retrieval, jointly score each pooled query–candidate pair with a cross-encoder that evaluates the query and candidate together to estimate their relevance, and use a supervised reranker to integrate the resulting evidence. We report Recall@5 and related top-K retrieval measures; we do not evaluate precision.

In this study we evaluate bi-encoders for source-to-CDE retrieval in caDSR, compare them with strong lexical baselines, and integrate them into a hybrid ranking system. We construct a benchmark of 69,102 expert-linked source-query/CDE pairs derived from caDSR metadata (68,142 after canonical deduplication and reachable-gold filtering). Retrieval is performed against a separate production catalog of 62,976 non-retired CDE records. We evaluate on an internal test set together with five distribution-shifted holdout sets designed to assess organizational and style-based shift. We study how query and CDE text representation, model choice, and two-stage fine-tuning with hard negatives affect retrieval quality.

Figure 1 provides an overview of the workflow, including benchmark construction, bi-encoder screening and fine-tuning, hybrid candidate generation, cross-encoder scoring, and supervised reranking. In the final system, FT-MPNet and CDE Match-Fuzzy contribute candidate CDEs to a shared candidate pool. For each query–candidate pair in this pool, a supervised histogram-based gradient-boosting classifier (HGBC) integrates features from the bi-encoder, keyword retriever, and FT-MedCPT cross-encoder to produce the final ranking.

**Figure 1:**
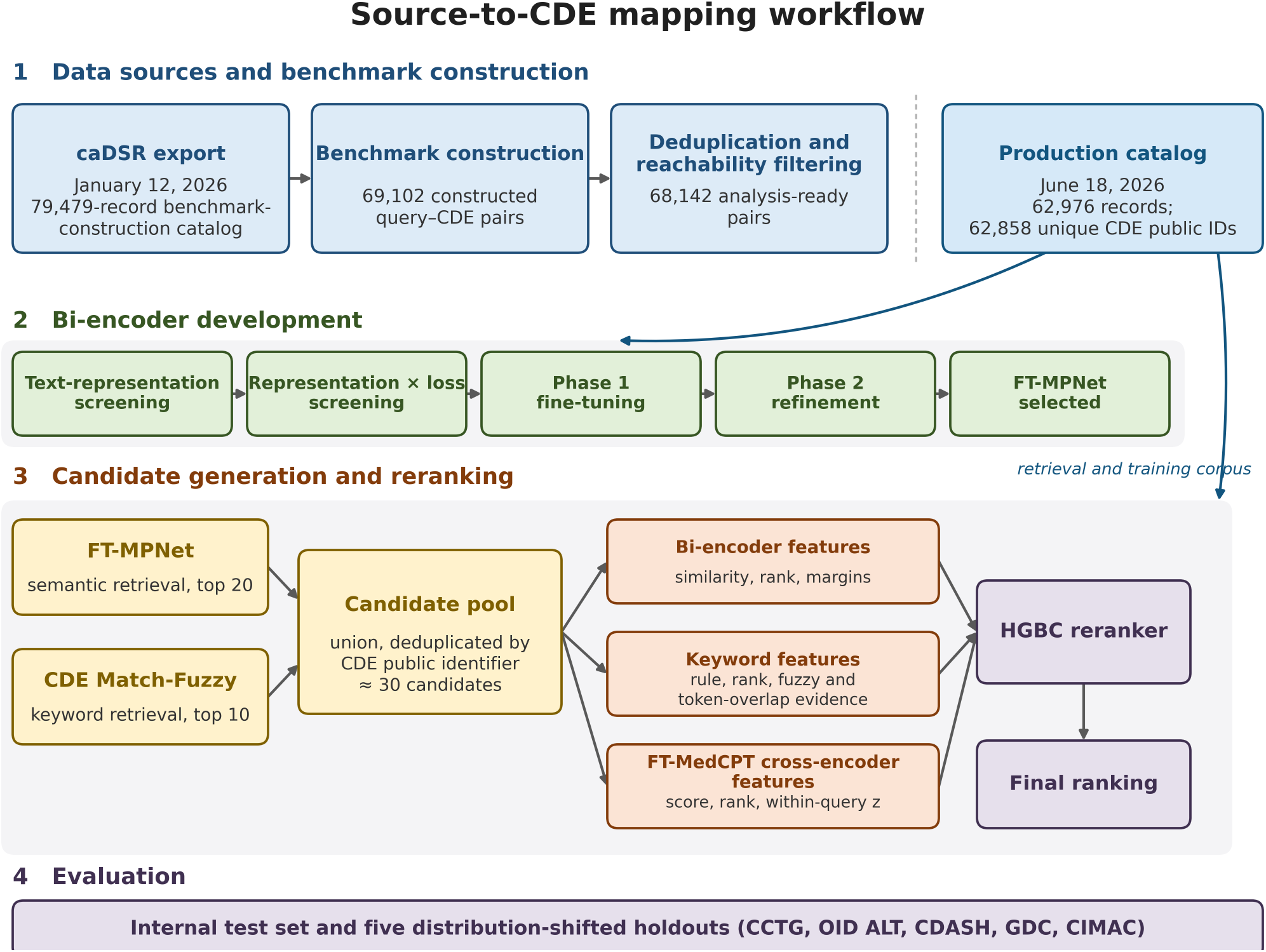
Source-to-CDE workflow. caDSR metadata yields benchmark queries, gold links, and splits; training and retrieval use a later production catalog. Bi-encoders are screened and fine-tuned, and FT-MPNet and CDE Match-Fuzzy form a candidate pool. Bi-encoder, keyword, and FT-MedCPT cross-encoder features are computed for each pooled candidate and supplied to the HGBC, which produces the final ranking. Evaluation uses the internal test set and five distribution-shifted holdouts.

The main contributions of this work are as follows:

1. A large benchmark dataset of expert-linked source-to-CDE mappings derived from caDSR metadata.
2. A systematic evaluation of bi-encoders, text representations, and fine-tuning strategies for source-to-caDSR retrieval.
3. A reproducible CDE Match-inspired keyword system, CDE Match-Fuzzy, that improves on the Python approximation to NCI CDE Match used for comparison.
4. A hybrid keyword and semantic retrieval system that outperforms the Python approximation to NCI CDE Match and all other evaluated methods on five of six datasets, while achieving near-ceiling performance on Genomic Data Commons (GDC) within one query of both the official service and its Python approximation.

## 2 Methods

### 2.1 Dataset, task, and evaluation sets

This study combines a caDSR-derived benchmark, three caDSR-derived distribution-shift holdouts, and two separately assembled external evaluation sets. The benchmark, the in-distribution train, validation and test splits, and the Canada Cancer Trials Group (CCTG), Object Identifier (OID) Alternate Name (ALT), and Clinical Data Acquisition Standards Harmonization (CDASH) holdouts were all constructed from the nightly caDSR XML export downloaded on January 12, 2026. The GDC and Cancer Immune Monitoring and Analysis Center (CIMAC) sets were assembled independently of that export and are used only as external evaluations. The retrieval task is to map a source-like query string to its correct CDE in caDSR. A bi-encoder independently encodes the source query and each candidate CDE description as vector embeddings. Candidates are ranked by cosine similarity, which measures the alignment between the query and CDE embedding vectors. Within the caDSR-derived benchmark and its holdouts, query strings were harvested from two metadata sources: alternate names (ALT), which are typically short label-like strings, and reference documents (REF) that serve as question or prompt text. To retain realistic source-side queries, extraction was restricted to selected curator-defined provenance sources, followed by normalization and deduplication within each CDE. Each query instance was linked to a stable CDE identifier. Additional extraction rules and provenance details are provided in the Supplementary Materials.

The caDSR-derived benchmark comprises 69,102 constructed query-CDE pairs, reduced to 68,659 after duplicate-pair removal. A query-level partition stratified by query source and provenance family gives the in-distribution train, Reranker Training, Validation and Test splits, totaling 64,346 pairs over 59,484 distinct queries. Three further caDSR-derived holdouts assess organizational and style shift and contribute 3,593 pairs: CCTG, OID ALT and CDASH. The independently assembled GDC and CIMAC evaluation sets contribute a further 203 pairs. Combining the in-distribution splits with all five holdouts, after filtering to queries whose gold CDE is present in the production catalog, gives an analysis-ready set of 68,142 pairs over 62,847 distinct queries. Benchmark construction used the January 2026 snapshot, whereas all fine-tuning and evaluation used a later production-oriented catalog of 62,976 CDE records. Exact split counts, catalog versions and filtering rules are reported in Supplementary Section S1.

### 2.2 Query and text representations

We evaluated multiple text representations for source queries and target CDEs. ALT and REF denote coarse query styles: ALT queries are short labels or identifiers, whereas REF queries are question- or prompt-like text. In the caDSR-derived datasets, these styles generally correspond to alternate-name and reference-document metadata, respectively; in independently assembled datasets, they were assigned by analogous query form.

For example, the ALT-style query CTP_CPS_prd_collect_dt_kn maps to *Product Collection Date Known Unknown Indicator*, whereas the REF-style query “Did patient require treatment with insulin therapy” maps to *Diabetes Mellitus Current Therapeutic Procedure Ind-2*, whose preferred question text is “Is patient being treated for diabetes.”

Query representations used these source-like strings, sometimes with concise permissible-value information. CDE representations combined metadata fields such as short name, long name, preferred definition, preferred question text, and permissible values, ranging from compact lexical forms to richer semantic descriptions.

Because permissible values can help distinguish enumerated elements, we also evaluated representations that included concise permissible-value summaries. True source-side permissible values were unavailable in the benchmark, so the query-side summary was a deliberately reduced, source-like rendering of the matched CDE’s permissible values, included only under predefined safeguards designed to limit leakage, whereas the CDE-side summary was a fuller rendering of the candidate CDE’s permissible values. Final representations were selected on the validation set; full recipes and construction rules are provided in the Supplementary Materials.

### 2.3 Models, baselines, and fine-tuning setup

This study evaluates three bi-encoders for source-to-CDE retrieval: the general-domain sentence-transformers/all-mpnet-base-v2 model (all-MPNet), the biomedical kamalkraj/BioSimCSE-BioLinkBERT-BASE model (BioSimCSE), and NeuML/pubmedbert-base-embeddings (PubMedBERT). Four comparators are reported. The official NCI CDE Match service is the production system described in the Introduction. For this study, official-service results are reported only for GDC and CIMAC because the other four evaluation sets were constructed from caDSR metadata fields searched directly by the live service, which could not be subjected to our query-level exact-match masking. The Python approximation to NCI CDE Match reimplements that service from its supplied PL/SQL source over the frozen production catalog, and unlike the live service it exposes its per-rule match types and can therefore carry the exact-match allowance described below. CDE Match-Fuzzy is our own CDE Match-inspired keyword retriever, which extends that matching flow with character- and word-level fuzzy and permissible-value evidence and supplies candidates to the final reranker. BM25 is a lexical baseline with no learned parameters; its weighting constants were left at standard values and only its query and CDE text representation was chosen on Validation, so it is deterministic and was run once. All retrieval methods used the same canonical query text and CDE catalog whenever their interfaces permitted it.

For semantic retrieval, query and CDE texts were encoded separately and ranked by cosine similarity. Each bi-encoder was evaluated in three stages: off-the-shelf inference, Phase 1 supervised contrastive fine-tuning with Multiple Negatives Ranking Loss (MNRL) or a symmetric bidirectional variant [21, 22], and Phase 2 refinement using mined hard negatives. Off-the-shelf screening compared query and CDE text representations, and the text representation and training objective carried forward into fine-tuning were selected separately for each model rather than shared across models. Fine-tuning used a standardized maximum sequence length for all models; Phase 1 used mixed-precision fine-tuning and Phase 2 was trained in FP32. All hyperparameters were selected on Validation, comparing configurations by the mean across two random seeds and prioritizing Recall@5, then Recall@10, and then MRR@100. A single seed was then retained from the winning seed-averaged configuration and used downstream. The internal test set and the distribution-shifted holdout sets were never used for tuning or model selection; they were evaluated only after validation-based selection was complete. Full training grids, loss-function screening, hard-negative strategies, and implementation details are provided in the Supplementary Materials.

FT-MPNet—all-MPNet [23] after the two-stage fine-tuning procedure described above—was selected as the final bi-encoder for downstream candidate generation and reranking. For each query, candidates retrieved by FT-MPNet were combined with candidates from the keyword retriever, and the union was deduplicated by CDE identifier to form the candidate pool passed to reranking. Bi-encoder, cross-encoder, and keyword features computed over this pool were supplied to a gradient-boosted reranker, which produced the final ranking.

Keyword retrieval used CDE Match-Fuzzy, an independently implemented, CDE Match-inspired method that combines exact, character- and word-level fuzzy, token-overlap and permissible-value evidence across five CDE metadata fields and contributes its top 10 candidates. It was selected over the Python approximation to NCI CDE Match on Validation. To limit circular exact lookup in the four caDSR-derived evaluation sets, both lexical methods used the same deterministic 70% query-level exact-match allowance; independently supplied GDC and CIMAC used 100%. Scoring and candidate-depth details are provided in Supplementary Section S5.2, and masking and sensitivity details are provided in Section S6.2.

Cross-encoder scoring evaluates each pooled query-candidate pair jointly. MiniLM [24], MedCPT and BGE [25] were compared off the shelf and after fine-tuning, and FT-MedCPT was selected by Validation Recall@5. Input representations, negative construction, sequence length and training details are provided in Supplementary Section S5.4.

## 3 Results

### 3.1 Descriptive dataset statistics

Table 1 summarizes the in-distribution splits and five holdouts. CCTG represents organizational REF-text shift, OID ALT compact identifier-style queries, CDASH standardized question text, GDC curated external metadata and CIMAC independently produced clinical-trial metadata.

**Table 1.** Dataset composition and evaluation-set roles. Query counts are reported after canonical deduplication and reachable-gold filtering. The ALT/REF mixture gives the share of ALT-style versus REF-style queries, as defined in Section 2.2.

| Dataset | Queries | ALT/REF mix | Shift summary |
| --- | --- | --- | --- |
| Train | 47,645 | ALT 40.5% / REF 59.5% | In-distribution caDSR pool (80% of query_ids); not filtered for whether the gold CDE is present in the production catalog |
| Reranker Training | 3,946 | ALT 40.6% / REF 59.4% | In-distribution; in-sample reference split (not used for model selection) |
| Validation | 3,934 | ALT 40.6% / REF 59.4% | In-distribution; primary model-selection split |
| Test | 3,959 | ALT 40.7% / REF 59.3% | same ALT/REF mixture as Train |
| CCTG | 1,097 | ALT 0.0% / REF 100.0% | Organizational shift: all REF-style question text from a held-out CCTG context |
| OID ALT | 1,766 | ALT 100.0% / REF 0.0% | Naming-style shift: all ALT-style short identifier-like queries |
| CDASH | 324 | ALT 0.0% / REF 100.0% | Standards-vocabulary shift:<br>REF-style CDISC CDASH<br>question text |
| GDC | 72 | ALT 45.8% / REF 54.2% | Curated external repository<br>shift: GDC<br>preferred-question-text +<br>alternate-name queries |
| CIMAC | 131 | ALT 100.0% / REF 0.0% | Real-world clinical-trial<br>submission shift: ALT-style<br>schema-like data-element<br>requests with PVs |

### 3.2 Text-representation heatmap screening

We screened 4 × 10 query-CDE representation grids for all-MPNet, BioSimCSE and PubMedBERT on Validation without fine-tuning. Because the representation preferred by off-the-shelf models may not remain optimal after fine-tuning, we carried forward two pairs for each model: a prespecified information-rich anchor that had performed well in early fine-tuning experiments, and the best-performing non-anchor pair from the off-the-shelf screen. These were compared directly in the subsequent loss-function screening. Figure 2 reports all-MPNet, for which the anchor ranked second and raw query text against preferred question text ranked first; biomedical-model heatmaps are in the Supplement.

**Figure 2:**
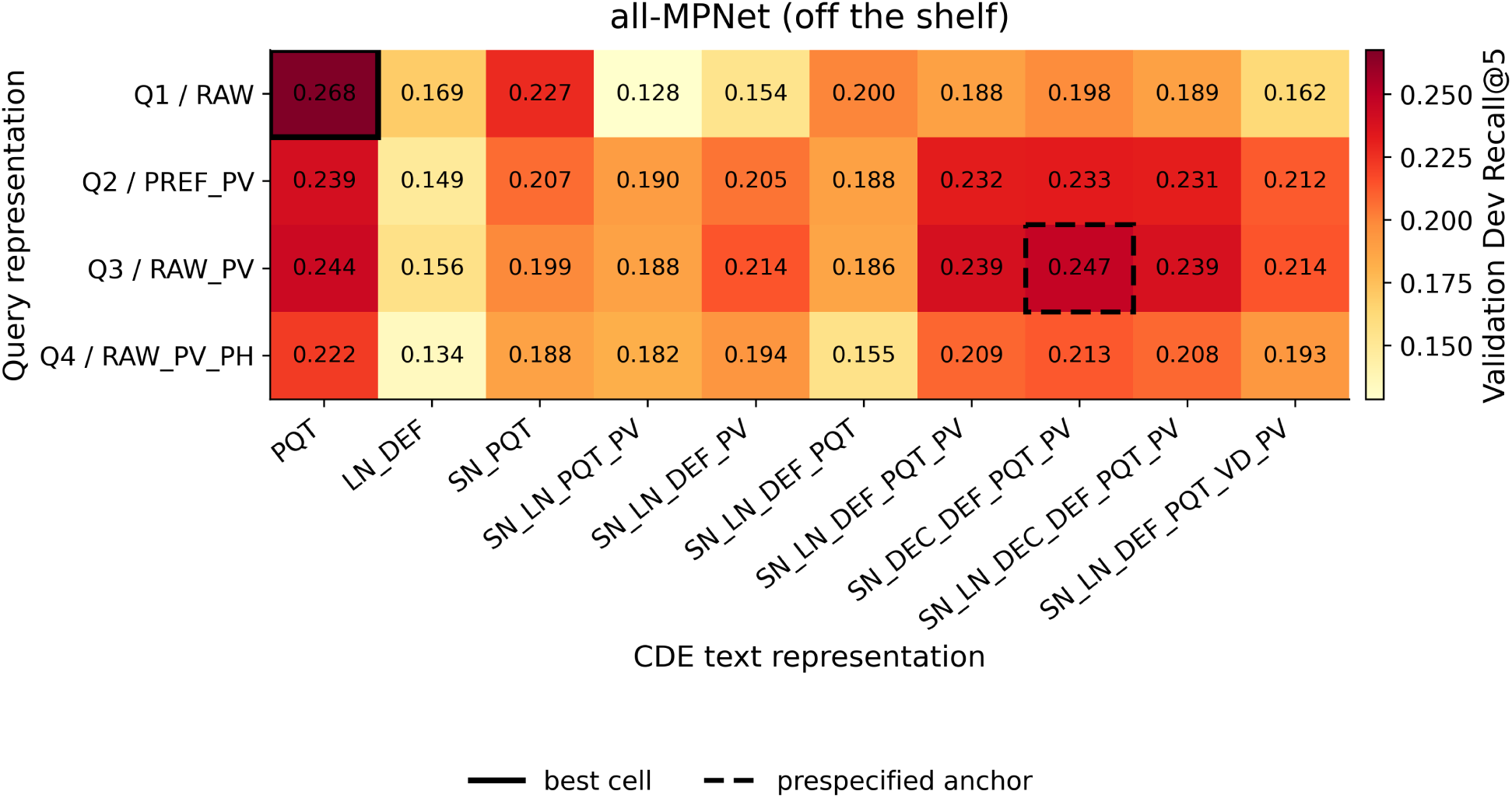
Off-the-shelf all-MPNet Validation Recall@5 across the 4 x 10 representation grid (n = 4,234 query–CDE pairs from 3,934 distinct Validation queries). The solid box marks the best cell and the dashed box the prespecified anchor.

### 3.3 Representation and loss-function screening

Because fine-tuning can change how effectively a bi-encoder uses a given text representation, we did not assume that the best off-the-shelf representation would remain best after training. For each bi-encoder, we therefore compared the prespecified anchor and the best off-the-shelf representation under Multiple Negatives Ranking Loss (MNRL) [21] and a symmetric bidirectional contrastive variant [22] before running the main hyperparameter sweep. Both loss variants were evaluated over two seeds. Tables 2 and 3 report the all-MPNet screen and the selected representation-loss combination for each model; the biomedical-model screens appear in Figure S3.

**Table 2.** all-MPNet representation-by-loss screen on Validation. Each cell is the mean of two seeds with the sample standard deviation; the selected condition is marked. The anchor pair is raw query + PV summary x SN + DEC + DEF + PQT + PV and the best pair is the highest-scoring non-anchor pair from Section 3.2.

| Representation pair | Query | CDE recipe | Loss | Val Recall@5 (mean) | SD |
| --- | --- | --- | --- | --- | --- |
| anchor | raw query + PV summary | SN + DEC + DEF + PQT + PV | MNRL | 0.8368 | 0.0023 |
| anchor | raw query + PV summary | SN + DEC + DEF + PQT + PV | Symmetric bidirectional | 0.8392 (selected) | 0.0013 |
| best | raw query | PQT | MNRL | 0.6436 | 0.0040 |
| best | raw query | PQT | Symmetric bidirectional | 0.6503 | 0.0042 |

**Table 3.**
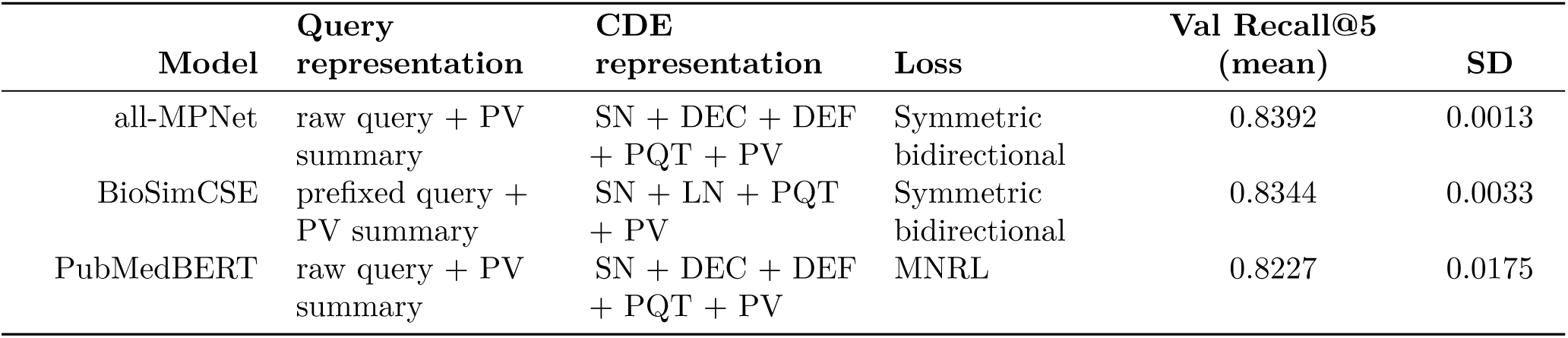
Selected query representation, CDE representation and loss for each retained bi-encoder, with the two-seed mean Validation Recall@5 and sample standard deviation from the representation-by-loss screen.

### 3.4 Bi-encoder selection on Validation

After selecting a representation and loss for each bi-encoder, we compared the models after the full two-stage fine-tuning procedure. Phase 2 improved all three bi-encoders (Figure 3). All three fine-tuned models achieved similar Validation performance, indicating that task-specific fine-tuning was effective across different embedding backbones. all-MPNet achieved the highest mean Validation Recall@5 (0.9014 +/- 0.0008), followed closely by PubMedBERT (0.8966) and BioSimCSE (0.8941), and was selected as FT-MPNet. The margins were small relative to two-seed variability and no paired statistical test was performed, so we do not claim statistical superiority.

**Figure 3:**
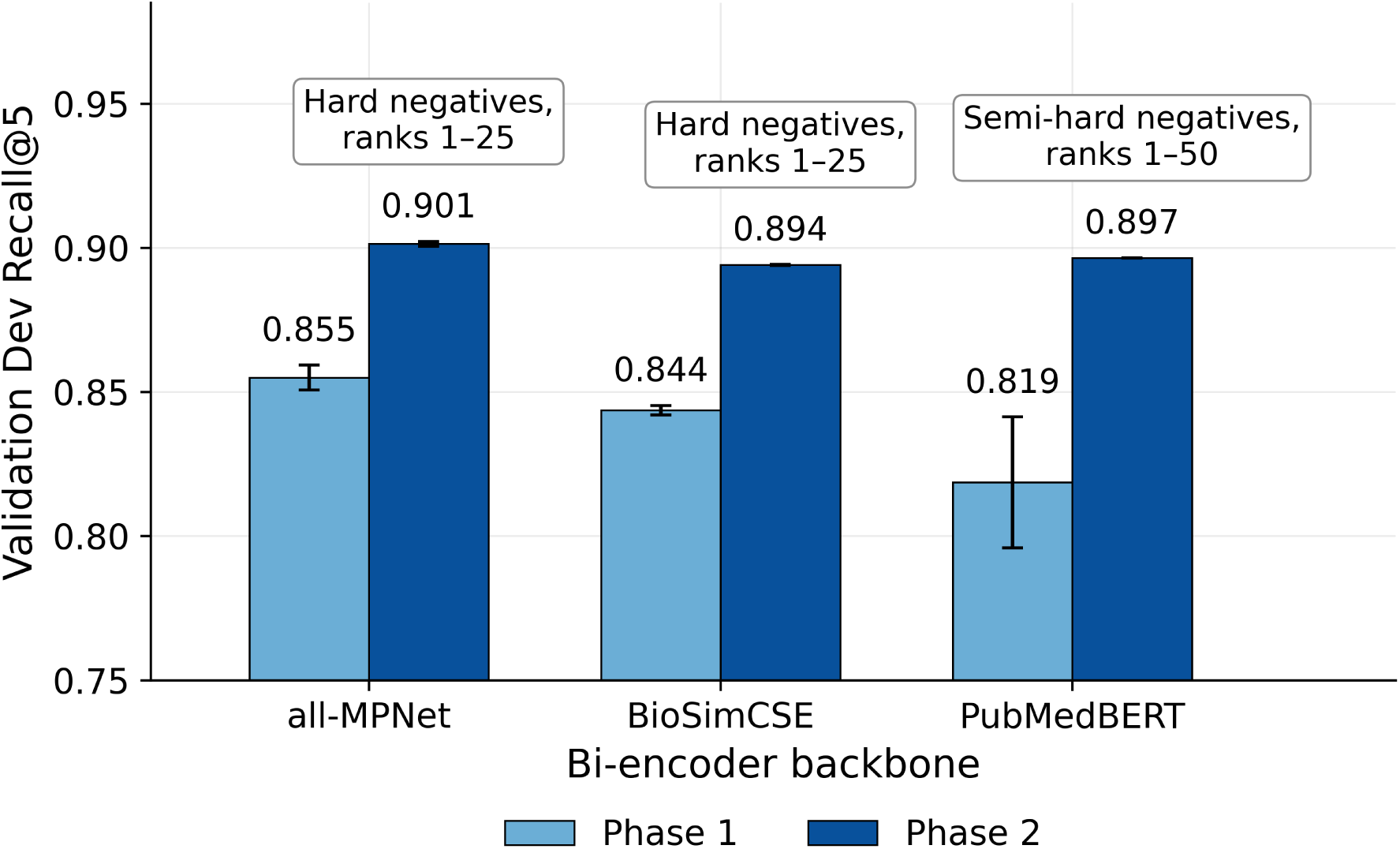
Validation Recall@5 after Phase 1 and Phase 2 fine-tuning (two-seed mean and sample SD). Selected Phase 2 strategies used hard negatives sampled from ranks 1–25 for all-MPNet and BioSimCSE and semi-hard negatives sampled from ranks 1–50 for PubMedBERT. Recall is computed over n = 4,234 query–CDE pairs from 3,934 distinct Validation queries.

Supplementary Section S4.5 summarizes performance across off-the-shelf, Phase 1, and Phase 2 stages for the three bi-encoders.

### 3.5 Cross-encoder selection on Validation

Three cross-encoder backbones were compared using Validation Recall@5: ncbi/MedCPT-Cross-Encoder, BAAI/bge-reranker-base, and cross-encoder/ms-marco-MiniLM-L-6-v2 (Figure 4). FT-MedCPT was selected for the final reranker. All three backbones used the prespecified anchor representation from the bi-encoder experiments, pairing the raw query text and its permissible-value summary with the SN + DEC + DEF + PQT + PV CDE recipe, and no separate text-representation search was performed for MedCPT, BGE or MiniLM.

**Figure 4:**
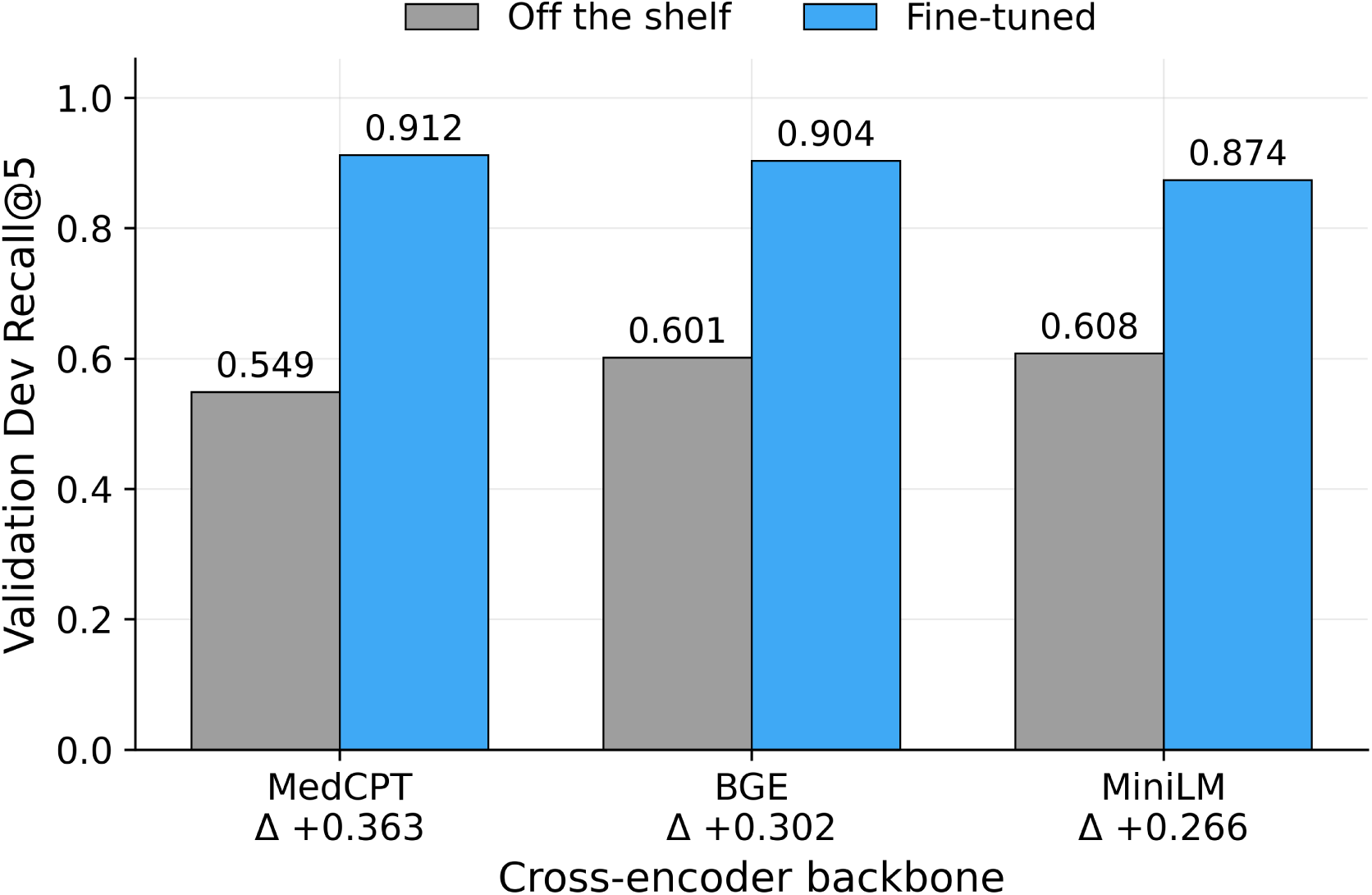
Off-the-shelf and fine-tuned cross-encoder Validation Recall@5. Δ denotes the difference between fine-tuned and off-the-shelf Recall@5. Each backbone used one fine-tuning run; FT-MedCPT was selected.

### 3.6 Keyword-algorithm selection and performance comparison

CDE Match-Fuzzy achieved Validation Recall@5 of 0.782 versus 0.728 for the Python approximation and was selected as the keyword retriever. Figure 5 reports full-set performance and performance among queries without available exact-match evidence for the gold CDE under the applicable allowance policy.

**Figure 5:**
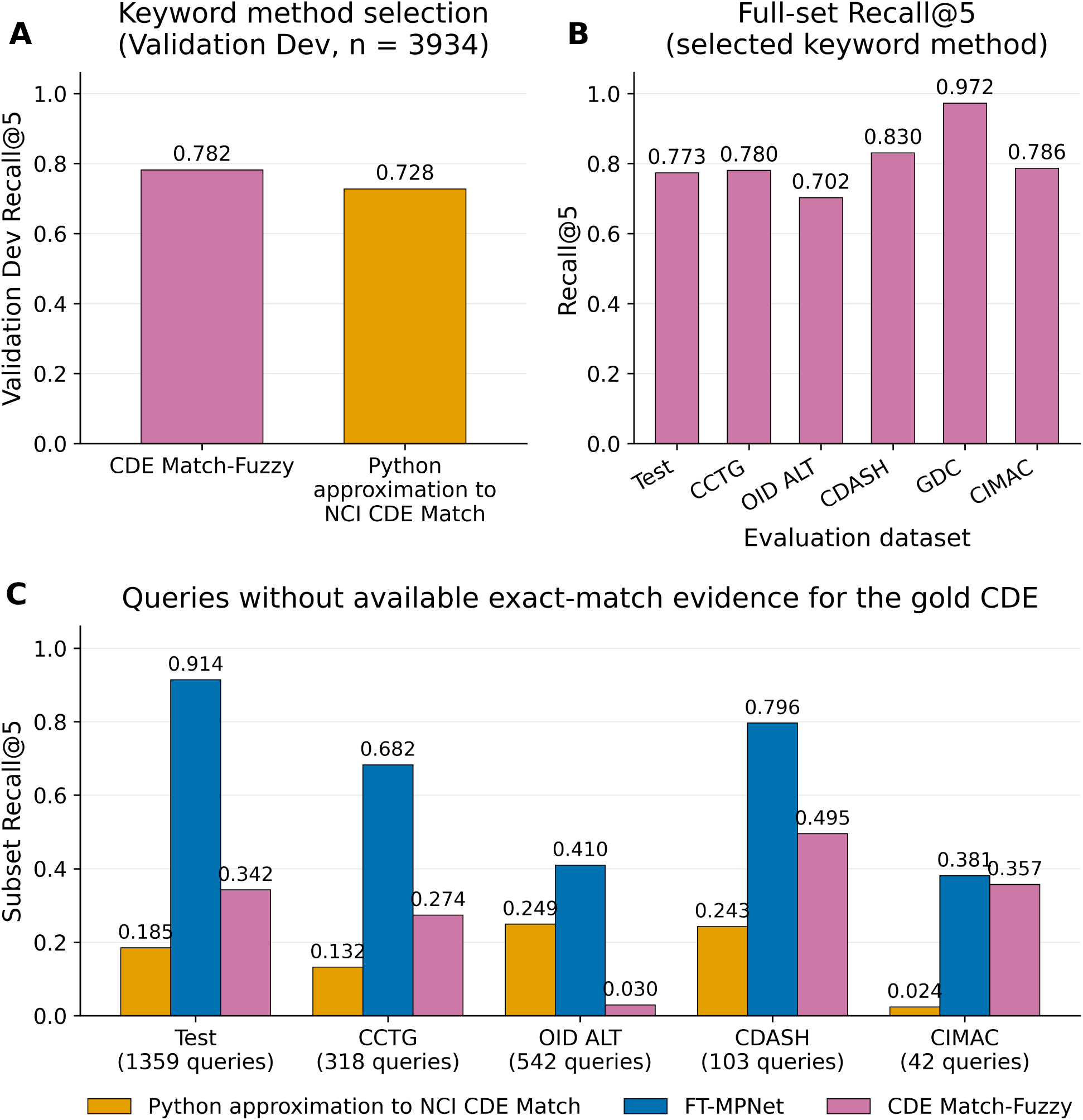
Keyword-method selection and evaluation: (A) Validation comparison; (B) CDE Match-Fuzzy Recall@5 across evaluation sets; and (C) Recall@5 among queries without available exact-match evidence for the gold CDE under the applicable allowance policy. GDC is omitted from panel C because only two such queries remain.

For this analysis, an exact lexical match means that the normalized query text is identical to at least one exact-match-eligible string for the gold CDE: its long name, a question-text string from its reference documents (including preferred or alternate question text) or an alternate name. Because the four caDSR-derived evaluation sets were constructed from metadata linked to their gold CDEs, unrestricted exact lookup could reveal the answer for many queries; exact matching to the gold CDE was therefore permitted for a deterministic 70% of queries. GDC and CIMAC were assembled independently and used a 100% allowance; the non-exact subset comprises queries for which no exact lexical match to the gold CDE was available under the applicable allowance policy.

### 3.7 End-to-end algorithm

The selected system combines FT-MPNet top-20 and CDE Match-Fuzzy top-10 candidates, deduplicates them by CDE public identifier, scores every pair with FT-MedCPT, and applies the HGBC reranker (Figure 6).

**Figure 6:**
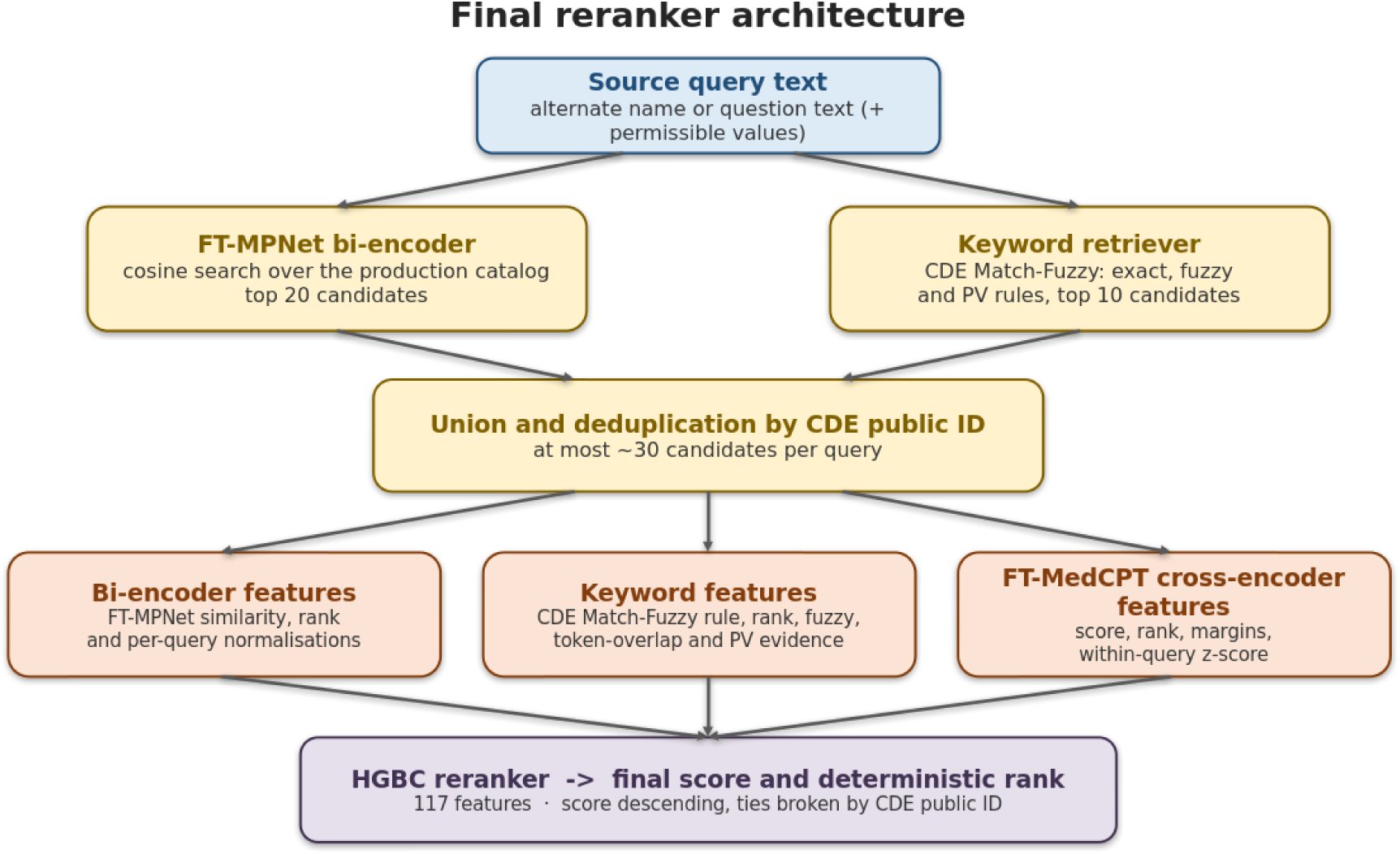
End-to-end reranker. FT-MPNet and CDE Match-Fuzzy form an approximately 30-candidate pool. For each candidate, the system computes bi-encoder, keyword, and FT-MedCPT cross-encoder features; the HGBC combines 117 lexical and semantic features to produce the final ranking. Ties are broken by CDE public identifier.

### 3.8 Final system performance

Table 4 summarizes Recall@5 for the final reranker and evaluated benchmarks.

**Table 4.** Recall@5 for the final reranker and evaluated benchmarks. FT-MedCPT reranks the same approximately 30-candidate pool used by the HGBC rather than retrieving from the full catalog. All tunable methods were selected on Validation.

| Dataset | Final reranker | Python approximation to NCI CDE Match | BM25 | FT-MPNet | CDE Match-Fuzzy | FT-MedCPT (top-30 pool reranking) |
| --- | --- | --- | --- | --- | --- | --- |
| Test | **0.971** | 0.719 | 0.361 | 0.912 | 0.773 | 0.914 |
| CCTG | **0.909** | 0.739 | 0.392 | 0.705 | 0.780 | 0.718 |
| OID ALT | **0.832** | 0.770 | 0.176 | 0.428 | 0.702 | 0.564 |
| CDASH | **0.920** | 0.750 | 0.710 | 0.840 | 0.830 | 0.864 |
| GDC | 0.972 | **0.986** | 0.542 | 0.833 | 0.972 | 0.903 |
| CIMAC | **0.802** | 0.679 | 0.160 | 0.565 | 0.786 | 0.595 |

The relative strengths of semantic and keyword retrieval varied across datasets, while the final hybrid reranker remained strong across these shifts. Together with the increased candidate coverage from combining the two retrievers (Table S5), this supports the value of integrating lexical and semantic evidence rather than relying on a single retrieval method.

At Stage 1, combining FT-MPNet and CDE Match-Fuzzy increased gold-CDE coverage over either retriever alone on every dataset (Table S5). A fixed-candidate-pool ablation further showed that lexical/keyword-oriented evidence contributed most to final ranking, suggesting that detailed lexical cues are especially useful for distinguishing among already-plausible CDE candidates (Figure S6).

The official NCI CDE Match service was evaluated only on GDC and CIMAC because it directly queries the caDSR alternate-name and question-text fields used to construct Test, CCTG, OID ALT, and CDASH. Evaluation on those four datasets would therefore be circular, and the live service could not be subjected to the query-level masking used in our controlled experiments. For CDE Match-Fuzzy and the Python approximation, exact matching to the gold CDE was permitted for a deterministic 70% of queries on the four caDSR-derived datasets and 100% on the independently assembled GDC and CIMAC sets. On GDC and CIMAC, both the official service and the Python approximation achieved Recall@5 of 0.986 and 0.679, respectively.

On GDC, the official service and Python approximation each retrieved the gold CDE for 71 of 72 queries, compared with 70 of 72 for the final reranker; the methods failed on different queries. On CIMAC, removing the 18 of 131 queries that recurred in training with the same gold CDE reduced Recall@5 from 0.802 to 0.779 for the final reranker and from 0.679 to 0.637 for the Python approximation, without changing method order (Table S6).

Exact-match allowance sensitivity analyses further showed that the final HGBC was comparatively robust to lexical exact-match availability (Figures S7 and S8). The fixed HGBC trained at the 70% allowance varied substantially less than either lexical method across the 0–100% allowance range.

### 3.9 Limitations

This study has several limitations. all-MPNet was selected on Validation, and the Phase 2 comparison rests on two seeds per configuration; the margins between the three fine-tuned bi-encoders are small (0.0048 Recall@5 over PubMedBERT and 0.0073 over BioSimCSE) and no paired statistical test was performed, so we make no claim that all-MPNet is significantly better than the biomedical models. On the six evaluation datasets the three fine-tuned bi-encoders are similarly close (Table S4), and the external evaluations are best read as evidence of robustness under distribution shift rather than as a ranking of model quality. The external sets are also not fully independent of training data under a text-plus-gold criterion (Table S6), and cross-encoder results come from a single training run per backbone, so they carry no seed variance. Finally, we report Recall@5 and related top-K measures; precision was not evaluated.

## 4 Discussion and Conclusion

Source-to-CDE mapping benefits from complementary evidence. Lexical matching resolves literal identifiers, abbreviations and short schema-like labels, where one rare token is decisive; dense semantic retrieval resolves the complementary case, in which a source element and its target CDE express the same concept in different words. Because the two families fail in different places, combining them is productive rather than redundant: on OID ALT, CDE Match-Fuzzy achieved Recall@5 of 0.702 versus 0.428 for FT-MPNet, whereas on the internal Test set FT-MPNet achieved 0.912 versus 0.773 for CDE Match-Fuzzy.

The later stages exploit that complementarity. The cross-encoder scores each query–candidate pair jointly, allowing it to model fine-grained interactions between specific query and CDE terms that independent bi-encoder embeddings may not fully preserve. The reranker then combines these interaction scores with lexical and semantic retrieval signals. When exact-match availability was varied from 0% to 100%, the HGBC trained at the 70% allowance maintained more stable Recall@5 than the lexical methods. Most of the remaining change in its performance arose because varying exact-match availability also changed whether the correct CDE entered the candidate pool.

Across the six evaluation datasets the integrated system achieved the highest Recall@5 among the evaluated methods on five. On GDC, near ceiling for lexical methods, direct CDE Match lookup retrieved one additional query of 72 and the systems failed on different queries; on CIMAC the integrated system was substantially stronger than the official service. These results indicate robust performance across heterogeneous datasets. This robustness matters because the final ranking is not tightly coupled to a particular level of exact-match availability. Overall, robust source-to-CDE mapping appears to depend not on any single retrieval signal, but on integrating complementary evidence within a unified ranking framework.

## Acknowledgements

Generative artificial intelligence tools were used in preparation of this work. Claude Code (Anthropic) and OpenAI GPT-5-series systems assisted with coding; ChatGPT (OpenAI GPT-5 series) assisted with manuscript revisions. Code produced with AI assistance was evaluated by the authors through targeted human review, numerical validation, and automated testing.

## Data Availability

Source code, reproducibility materials, and the processed GDC and CIMAC evaluation datasets used in this study are publicly available in the DEMap repository at https://github.com/jhtanis/DEMap. The CIMAC evaluation set was derived from the CIMAC-CIDC clinical data-element template, available from the National Cancer Institute at https://dctd.cancer.gov/research/networks/precision-medicine-oncology/cimac-cidc. The caDSR-derived benchmark and production catalog were constructed from the January 12, 2026 and June 18, 2026 caDSR CDE XML exports, respectively; these dated exports are available from the caDSR CDE XML Archive at https://cadsr.nci.nih.gov/ftp/caDSR_Downloads/CDE/XML_Archive/. Additional provenance and dataset-construction details are provided in the repository documentation and Supplementary Materials.

## Supplementary Materials

### S1. Data sources, benchmark construction, and evaluation sets

#### S1.1 Data sources and retrieval task

This study uses three data resources: a caDSR-derived benchmark and three caDSR-derived holdouts constructed from the January 12, 2026 XML export; independently assembled GDC and CIMAC evaluation sets; and a production-oriented CDE catalog derived from the June 18, 2026 caDSR export. The January export supplied the source-like queries and gold CDE links used for benchmark construction, whereas the June catalog supplied the candidate CDEs used for training and retrieval.

The task is to map a source-like query string to its accepted caDSR Common Data Element (CDE). Retrieval methods rank candidate CDEs from the production catalog, and a query is counted as successfully retrieved when an accepted gold CDE appears within the evaluated rank cutoff.

#### S1.2 Query construction and provenance

The January export was parsed into normalized tables containing CDE metadata, alternate names, reference documents, and permissible values. Queries were derived from alternate names (ALT), which are typically short labels or identifier-style strings, and reference documents (REF), which are typically questions or prompt text.

Extraction was restricted to curator-defined provenance sources. ALT queries were selected using specified combinations of alternate-name type and context, together with deterministic concatenation rules where applicable. REF queries were selected from question-like document types, including Preferred Question Text, Alternate Question Text, and Application Standard Question Text. Queries were normalized, deduplicated within each CDE and provenance group, and linked to stable CDE public identifiers.

ALT and REF query styles are defined in Section 2.2. For the caDSR-derived datasets, these styles correspond to queries harvested from alternate-name and reference-document metadata, respectively. Provenance family is a separate curator-harvest grouping used for stratification and holdout construction; a provenance family can contain both ALT- and REF-style queries.

#### S1.3 Benchmark splits and evaluation sets

The benchmark contains 69,102 constructed query–CDE pairs. After canonical deduplication and filtering to queries whose gold CDE was present in the production catalog, 68,142 pairs remained, representing 62,847 distinct queries.

The in-distribution data comprised 64,346 pairs representing 59,484 distinct queries. These were partitioned at the query level into Train, Reranker Training, Validation, and Test, with stratification by query source and provenance family. Approximately 80% of the queries were assigned to Train, with the remainder divided among the three evaluation splits.

Five additional holdouts were used to evaluate distribution shift. CCTG, OID ALT, and CDASH were derived from the January caDSR export but separated by provenance or query style. GDC and CIMAC were assembled independently of that export and used only for external evaluation. Final dataset sizes are reported in Table 1.

#### S1.4 CDE catalogs and reachability filtering

Two caDSR catalogs served different purposes. The January benchmark-construction catalog contains 79,479 records representing 78,533 unique CDE public identifiers. It was not restricted to production-eligible records and included 15,852 records with a retired workflow status, together with a small number of records from administrative contexts. This catalog defined the source metadata and gold CDE links used to construct the caDSR-derived queries.

The June production catalog contains 62,976 records representing 62,858 unique CDE public identifiers. It was created from a 79,827-record export by removing 16,846 retired records and 21 records from administrative TEST or Training contexts; 16 records met both exclusion criteria. This filtering was administrative and unrelated to the machine-learning data splits.

The production catalog supplied CDE text for training pairs, in-batch negatives, candidate generation, and evaluation. Evaluation sets were filtered to retain only queries whose accepted gold CDE was present in this catalog.

Because the catalog was filtered and derived from a later export, 622 training pairs lacked target CDE text and were removed, leaving 50,972 training pairs.

#### S1.5 Training–evaluation overlap

We assessed whether source-to-CDE mappings in the five holdouts duplicated mappings in Train. We first compared query identifiers. CCTG, OID ALT, GDC, and CIMAC shared no query identifiers with Train. CDASH shared 14 identifiers with the in-distribution data, but the associated accepted gold CDE sets were disjoint; these records therefore did not represent duplicate mappings and were retained.

We then applied a stricter, content-based definition: identical normalized query text paired with the same gold CDE public identifier. Under this definition, repeated mappings occurred in 8 of 1,097 CCTG queries, 7 of 1,766 OID ALT queries, 14 of 324 CDASH queries, 4 of 72 GDC queries, and 18 of 131 CIMAC queries. Table S6 reports sensitivity analyses after removing these repeated mappings, and leakage-filtered evaluation files are provided alongside the original sets.

### S2. Dataset characterization

#### S2.1 ALT/REF style and provenance composition

**Table S1.**
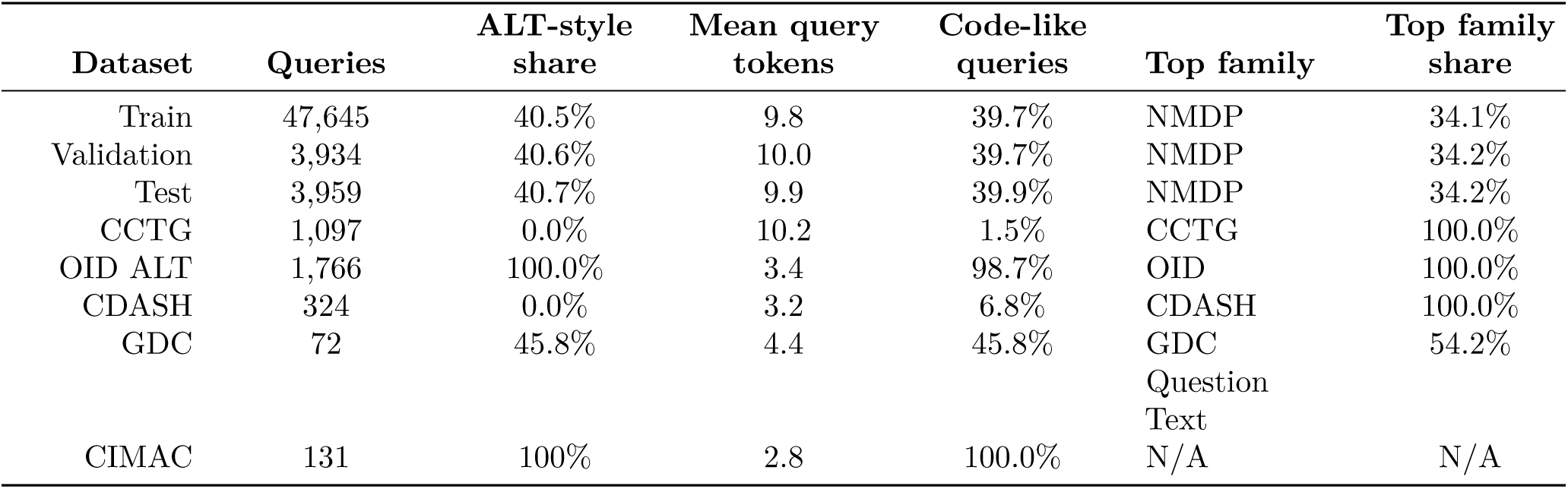
ALT/REF-style and provenance composition of the training and evaluation datasets.

- **Queries:** number of queries after canonical deduplication and reachable-gold filtering.
- **ALT-style share:** share of unique queries classified as ALT style rather than REF style, using the definitions in Section 2.2.
- **Mean query tokens:** mean number of tokens per query, where a token is a maximal run of [A-Za-z0-9] characters in the bare source string (no ALT_NAME:/REF_TEXT: label and no permissible-value block).
- **Code-like queries:** share of queries that are a single schema-style identifier with no internal whitespace.
- **Top family:** most common provenance family in that dataset.
- **Top family share:** share of queries belonging to that provenance family.

Train, Validation, and Test were constructed to have similar ALT/REF-style and provenance profiles. The holdouts intentionally differ in style: CCTG and CDASH contain REF-style question text, OID ALT and CIMAC contain ALT-style identifier-like queries, and GDC combines both styles. The provenance-family columns separately describe the curator or external-source groupings represented in each dataset. For CIMAC, these fields are N/A because the evaluation set spans multiple clinical trials rather than coming from known provenance families.

#### S2.2 Query-style composition

**Table S2.**
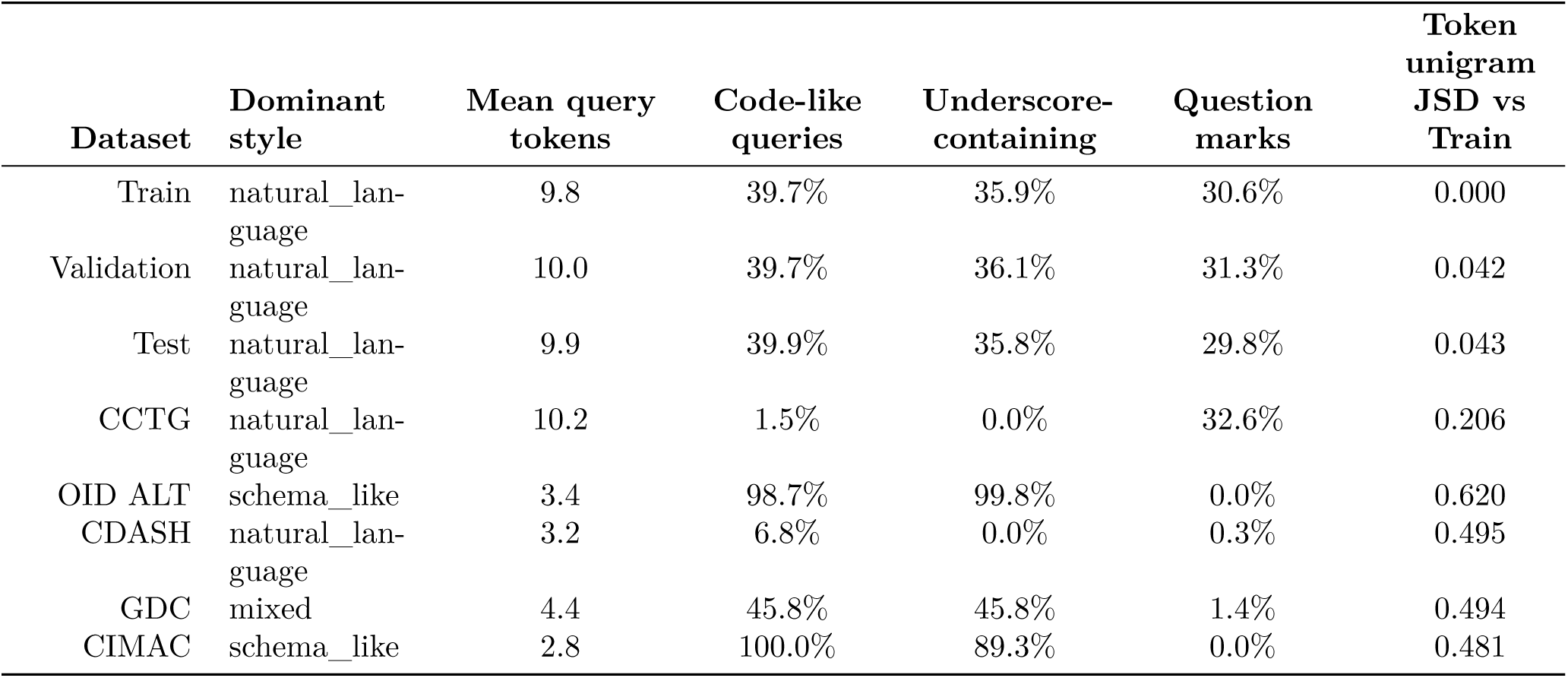
Query-style composition of the training and evaluation datasets.

- **Dominant style:** coarse style label assigned to the dataset (natural_language, schema_like or mixed).
- **Mean query tokens:** mean number of tokens per query, a token being a maximal run of [A-Za-z0-9] characters in the bare source string.
- **Code-like queries:** share of queries that are a single schema-style identifier with no internal whitespace.
- **Underscore-containing:** share of queries containing at least one underscore.
- **Question marks:** share of queries containing at least one question mark.
- **Token unigram JSD vs Train:** Jensen-Shannon divergence (natural logarithm, additive smoothing 1e-8) between the dataset’s token unigram distribution and the Train reference distribution; larger values indicate a stronger lexical shift away from training.

Higher token-distribution divergence indicates a stronger shift away from the training query distribution. OID ALT and CIMAC are schema-like: nearly every query is a single identifier, underscores are near-universal and question marks are absent. CDASH is short natural language rather than schema text. Divergence from Train increases with distance from the training distribution, in the order Validation 0.042 < Test 0.043 < CCTG 0.206 < CIMAC 0.481 < GDC 0.494 < CDASH 0.495 < OID ALT 0.620.

### S3. Text-representation design and screening

#### S3.1 CDE metadata fields

CDE text representations were constructed from fields available in the caDSR XML export: preferred name (abbreviated here as short name, SN), long name (LN), preferred definition (DEF), data element concept long name (DEC), preferred question text (PQT), value domain (VD), and permissible-value summary (PV). SN is the closest lexical analogue of a source variable name, whereas LN, DEF, DEC, and PQT provide additional semantic context. PV summaries were included only for enumerated CDEs.

#### S3.2 Permissible-value summaries

When permissible values were available, we constructed PV summaries for the CDE side and, in selected cases, for the query side. Because true source-side permissible values were unavailable, both summaries were derived from the permissible-value inventory of the matched CDE. Query-side PV summaries were included only when the case was not judged to be leakage-prone.

To reduce leakage, query-side PV summaries were designed to be more conservative and more source-like than CDE-side summaries. The query side used a lower display cap, allowed mild preference for generic concepts such as yes/no, and favored code-oriented rendering when available. The CDE side used preferred meaning labels and a higher display cap, yielding a more formal representation. Additional safeguards omitted selected small PV sets on the query side, retried deterministic sampling when rendered query and CDE summaries were identical, and suppressed cases in which the query-side summary was a strict subset of the CDE-side summary.

#### S3.3 Query text variants

We evaluated four query representations:

- Q1: raw query text only
- Q2: prefixed query text, with PV summary when available
- Q3: raw query text plus PV summary when available
- Q4: raw query text plus PV summary when available, with an explicit missing-value block when PV summary was absent

Q1 is the simplest source-like representation. Q2 tests whether explicit labels for query content and answer-choice information improve retrieval. Q3 removes those labels to test whether the extra structure in Q2 is helpful. Q4 further tests whether explicitly marking missing PV information is more helpful than leaving it absent. Together, these variants assess whether answer-choice information and missingness cues improve source-to-CDE matching.

For independently assembled datasets, Q2 prefixes were assigned from the ALT/REF query-style classification rather than from caDSR metadata provenance.

#### S3.4 CDE text recipes

The ten CDE text recipes evaluated in the representation grid, named by the fields they concatenate, are: PQT; LN + DEF; SN + PQT; SN + LN + DEF + PQT; SN + LN + DEF + PV; SN + LN + PQT + PV; SN + LN + DEF + PQT + PV; SN + DEC + DEF + PQT + PV; SN + LN + DEF + PQT + VD + PV; SN + LN + DEC + DEF + PQT + PV. Fields are joined in the order listed with a “ | ” separator and a field label prefix.

SN values were omitted when they were purely numeric or version-like strings, since these were unlikely to aid semantic matching. For non-enumerated CDEs, the PV component was omitted.

#### S3.5 Permissible-value overlap analysis

**Table S3.** PV-summary overlap between query-side and CDE-side permissible values.

| Group | Rows with<br>query+CDE<br>PV | Normalized<br>block match | Query shorter<br>than CDE | Mean Jaccard<br>Overlap |
| --- | --- | --- | --- | --- |
| overall | 39,391 | 0.4% | 30.1% | 0.168 |
| ENUM | 24,543 | 0.4% | 45.2% | 0.153 |
| BINARY_WITH_UN-KNOWN | 3,074 | 0.1% | 0.0% | 0.333 |
| BINARY_WITH_NA | 8,816 | 0.0% | 0.0% | 0.123 |

- **Group:** PV-type subgroup used for the overlap analysis, such as overall, ENUM, or binary-family types. Rows with query+CDE PV: number of rows where both the query and the matched CDE have PV summaries present.
- **Normalized block match:** percentage of rows where the full query-side PV block matches the CDE-side PV block after normalization.
- **Query shorter than CDE:** percentage of rows where the query-side PV summary contains fewer items than the CDE-side PV summary.
- **Mean Jaccard Overlap:** mean Jaccard overlap between the query-side and CDE-side permissible-value item sets.

Whole-block agreement between query-side and CDE-side permissible values is rare: over the 39,391 rows in which both sides carry a permissible-value summary, only 0.4% match as a normalized block, while the mean item-level Jaccard overlap is 0.168 and 30.1% of query-side summaries are shorter than the CDE-side summary. Query-side permissible values therefore contribute partial, not duplicated, evidence.

#### S3.6 Off-the-shelf representation screening

**Figure S1:**
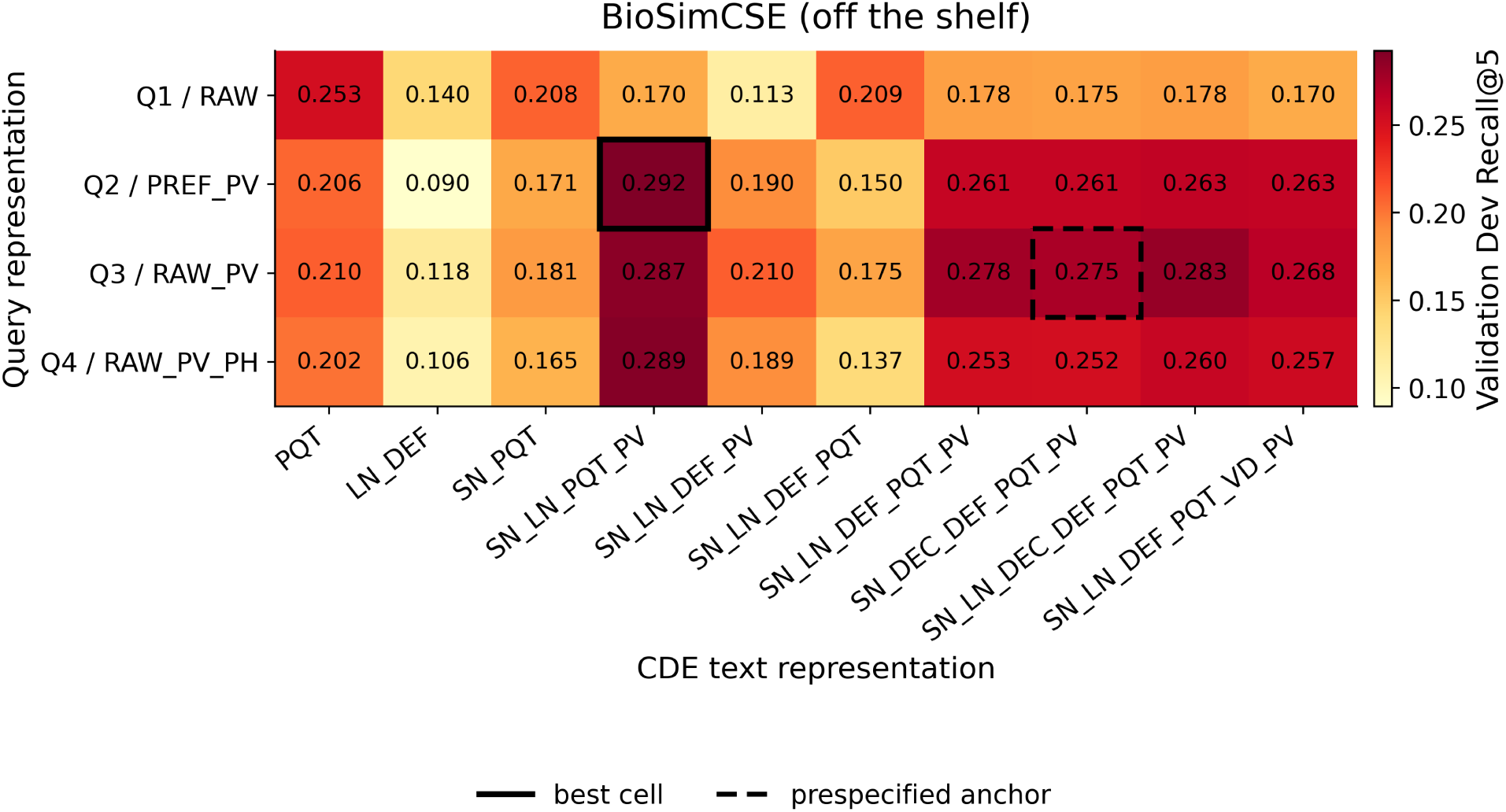
Validation Recall@5 across the 4 x 10 query-CDE text-representation grid for off-the-shelf BioSimCSE (n = 4,234 query–CDE pairs from 3,934 distinct Validation queries). The solid box marks the best cell and the dashed box the prespecified anchor.

We screened the final 40 query-CDE representation combinations, formed from four query variants and ten CDE recipes, separately for all-MPNet, BioSimCSE, and PubMedBERT. For each model, the prespecified SN + DEC + DEF + PQT + PV anchor and the best-performing non-anchor representation pair on Validation were carried forward to the 2 x 2 representation and loss-function screen. This screening stage used off-the-shelf inference at each model’s own effective maximum sequence length: 384 tokens for all-MPNet, 256 tokens for BioSimCSE (whose 128-token checkpoint value is raised to the model loader’s 256-token floor), and 512 tokens for PubMedBERT. Fine-tuning subsequently standardized the maximum sequence length across all models, so these off-the-shelf values apply only to this stage.

### S4. Bi-encoder development and evaluation

#### S4.1 Bi-encoder models, retrieval setup, and selection procedure

We evaluated three bi-encoders. The general-domain model was sentence-transformers/all-mpnet-base-v2 (all-MPNet), a general-purpose bi-encoder based on MPNet.

The biomedical models were kamalkraj/BioSimCSE-BioLinkBERT-BASE (BioSimCSE) and NeuML/pubmedbert-base-embeddings (PubMedBERT). BioSimCSE builds on BioLinkBERT [26] and was developed for biomedical sentence similarity and retrieval [12]. PubMedBERT provides domain-specific biomedical language representations [11] and the NeuML checkpoint adapts it for sentence-level semantic search and similarity.

For bi-encoder retrieval, query and CDE texts were encoded separately and candidate CDEs were ranked by cosine similarity. Model and text-representation selection were performed on the Validation split, using Recall@5 as the primary criterion and Recall@10 and MRR@100 as secondary ranking-sensitive criteria. The internal test set and the distribution-shifted holdout sets were never used for tuning, model selection, or checkpoint selection; they were evaluated only after validation-based selection was complete, using the final selected version of each model.

The same selection procedure was applied at Phase 1 and Phase 2. Each complete hyperparameter configuration was run with two seeds (0 and 1) and the two runs were averaged. Configurations were ordered first by mean Validation Recall@5; Recall@10 and then MRR@100 were consulted only to break exact ties in the preceding metric. Every run trained for a fixed number of epochs and used the resulting final checkpoint; no within-run checkpoint selection was performed. A single seed from the winning configuration supplied the checkpoint used downstream.

**Figure S2:**
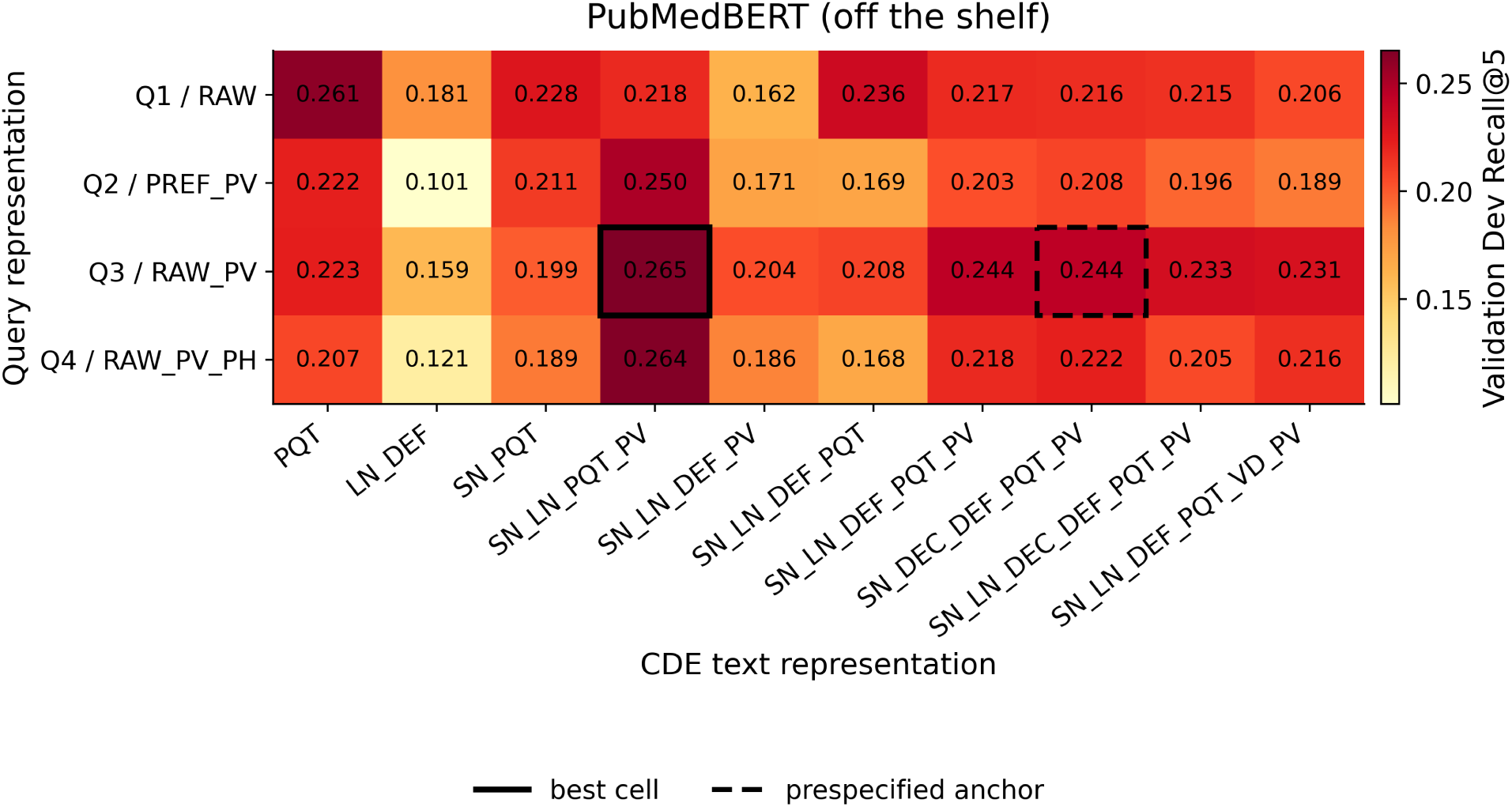
Validation Recall@5 across the same 4 × 10 query–CDE text-representation grid for off-the-shelf PubMedBERT (n = 4,234 query–CDE pairs from 3,934 distinct Validation queries). The solid box marks the best cell and the dashed box the prespecified anchor.

#### S4.2 Representation-by-loss screening

After fixing text-representation and formatting choices, we screened, for each model, the prespecified anchor representation pair and that model’s best-performing non-anchor pair from the off-the-shelf screen, under MNRL [21] and a symmetric bidirectional contrastive variant [22]. Each screening run used 3 epochs, learning rate 5 *×* 10*^−^*^5^, temperature 0.05, and two random seeds. The best-performing representation-loss combination was then selected independently for each model and used in that model’s main Phase 1 sweep.

#### S4.3 Phase 1 fine-tuning

The Phase 1 hyperparameter grid was: learning rate {7 *×* 10*^−^*^5^, 1 *×* 10*^−^*^4^, 1.5 *×* 10*^−^*^4^}, temperature {0.04, 0.07, 0.10}, and epochs {1, 2, 3}, giving 27 configurations per model and, across two seeds, 54 runs per model. Both the representation-and-loss screen and the Phase 1 sweep used a maximum sequence length of 256 tokens, a physical batch size of 128 query-positive pairs - at most 256 encoded strings per optimizer step - and gradient accumulation of 1. Phase 1 executed with float16 autocast, FP32 model parameters, and gradient scaling. To reduce false negatives during in-batch contrastive training, we used a no-duplicate batch sampler that prevented repeated normalized query text or repeated normalized CDE text within the same batch.

At the 256-token fine-tuning limit, query truncation was negligible: only one of the 55,525 distinct fine-tuning queries exceeded the limit for all-MPNet, and none did for BioSimCSE or PubMedBERT. On the CDE side, 940 of 62,976 catalog records (1.49%) exceeded 256 tokens under the SN + DEC + DEF + PQT + PV representation.

#### S4.4 Phase 2 hard-negative refinement

Phase 2 began from the selected Phase 1 checkpoint for each model and used the same loss family selected in Phase 1. Each configuration was trained for one epoch with two random seeds, a maximum sequence length of 256 tokens, gradient accumulation of 1, and FP32 parameters without autocast or gradient scaling.

To make the negative-mining strategies comparable, each optimizer step encoded 132 text sequences. The condition with no mined negatives used 66 anchor–positive pairs, whereas the mined-negative conditions used 11 training examples, each containing one anchor, one positive, and 10 negatives. Learning-rate and temperature grids were defined relative to the corresponding Phase 1 winner:

**Figure S3:**
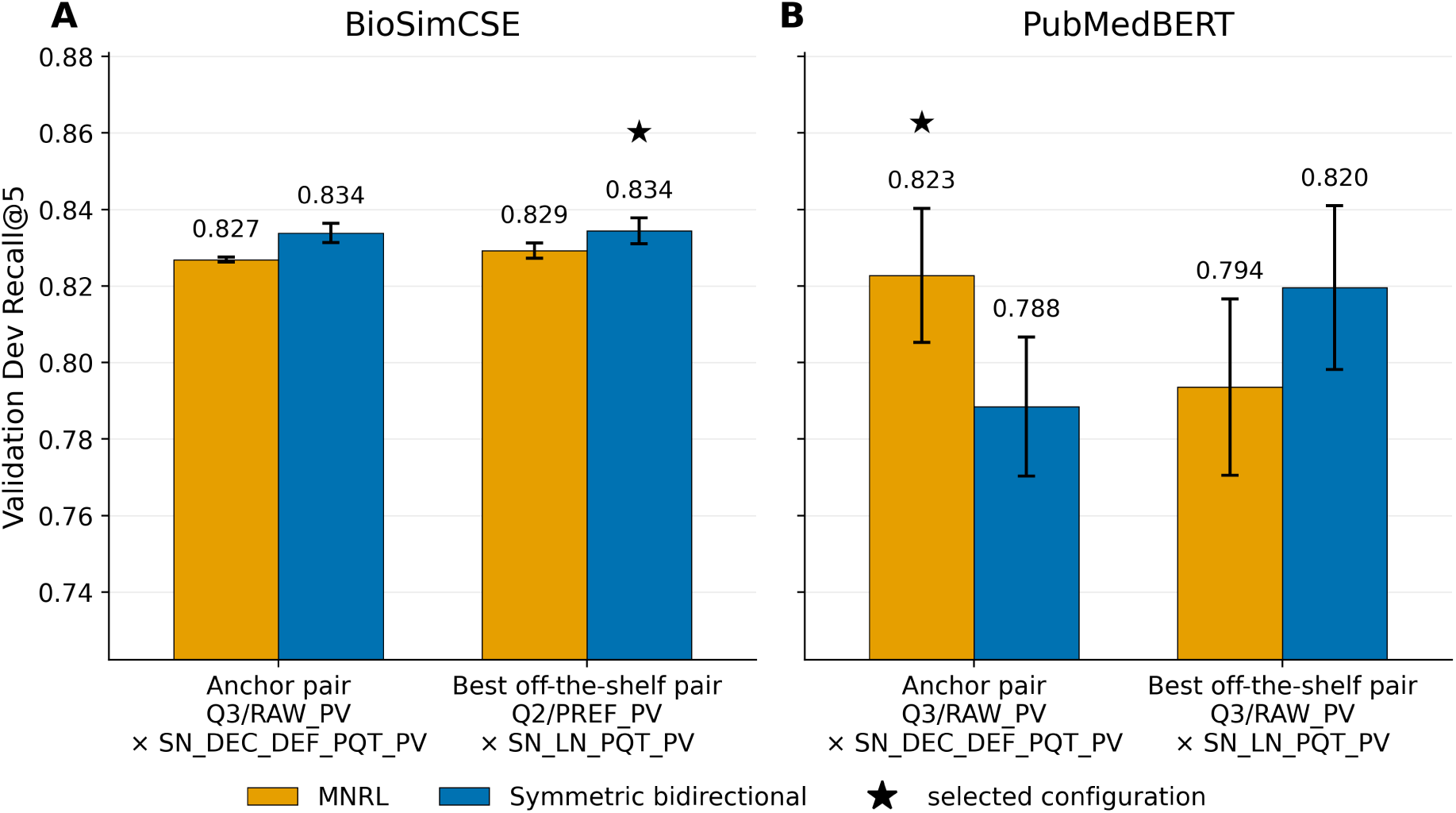
Validation Recall@5 for BioSimCSE (left) and PubMedBERT (right) across the two representation pairs and the MNRL and symmetric bidirectional losses, on a shared scale. Bars show the two-seed mean and error bars the sample standard deviation; the selected condition is marked.

- 7 *×* 10*^−^*^5^ *→ {*3 *×* 10*^−^*^5^, 5 *×* 10*^−^*^5^, 7 *×* 10*^−^*^5^*}*
- 1 *×* 10*^−^*^4^ *→ {*5 *×* 10*^−^*^5^, 7 *×* 10*^−^*^5^, 1 *×* 10*^−^*^4^*}*
- 1.5 *×* 10*^−^*^4^ *→ {*7 *×* 10*^−^*^5^, 1 *×* 10*^−^*^4^, 1.5 *×* 10*^−^*^4^*}*

The temperature mappings were:

- 0.04 → {0.03, 0.04, 0.05}
- 0.07 → {0.04, 0.06, 0.08}
- 0.10 → {0.07, 0.08, 0.10}

We compared three strategies: no mined negatives, hard negatives sampled from ranks 1–25, and semi-hard negatives sampled from ranks 1–50. The negative-mining strategy was treated as part of the hyperparameter configuration, and configurations were selected using the same Validation procedure described in Section S4.1.

FT-MPNet was selected as the final bi-encoder for downstream candidate generation and reranking. Where Phase 2 results for the biomedical bi-encoders are reported, they are descriptive; differences of the observed magnitude fall within seed-to-seed variability and are not interpreted as evidence of statistical superiority.

#### S4.5 Bi-encoder performance

Figure S4 summarizes Validation performance across off-the-shelf inference, Phase 1 fine-tuning, and Phase 2 refinement. Table S4 reports the retained Phase 2 models on the six evaluation datasets and includes their Validation selection results for reference.

**Table S4.**
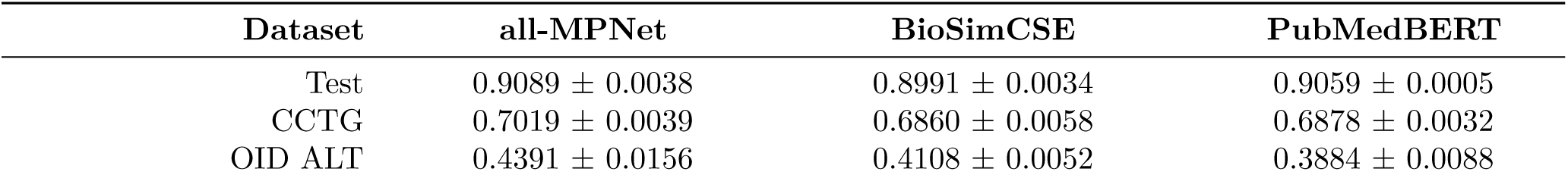
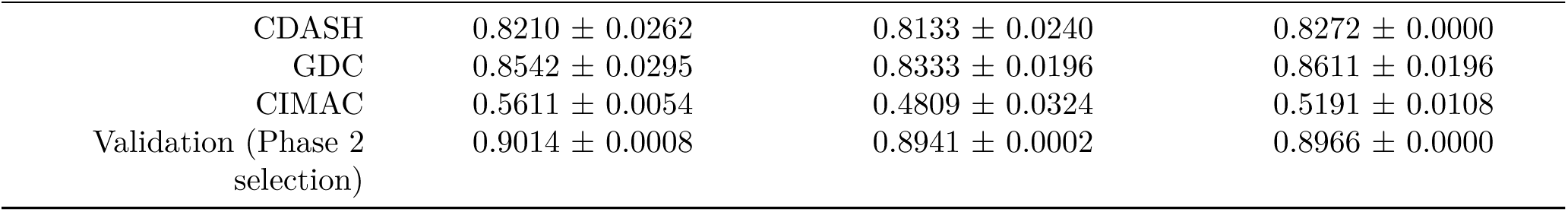
Bi-encoder Recall@5 on the six canonical evaluation datasets. Each cell is the mean over the two seeds of that model’s winning Phase 2 configuration, with the sample standard deviation; evaluation is inference-only on the frozen checkpoints, query-level against reachable gold, with public-identifier deduplication, over the same June production catalog (62,976 CDEs).

For CIMAC, which was assembled independently of caDSR, the Q2 prefix reflects its ALT-style classification (Section 2.2) rather than caDSR metadata provenance.

### S5. Baselines and hybrid reranking system

#### S5.1 BM25 baseline

BM25 was Okapi BM25 with the standard Lucene non-negative inverse-document-frequency formulation, computed over unigram term counts. The weighting constants were left at their conventional values of k1 = 1.2 and b = 0.75 and were never tuned; query terms were weighted binarily, so a repeated query term contributes once. Documents and queries were tokenized identically: Unicode normalization, case folding, and splitting on runs of alphanumeric and underscore characters. No stop-word list, stemming, lemmatization or n-gram expansion was applied, and no vocabulary pruning was used. Fields absent for a given CDE were omitted rather than replaced by a placeholder.

BM25 retrieved from the same production CDE catalog as the neural methods and was evaluated on the same six canonical datasets with the same reachable-gold filtering, query-level denominators, and public-identifier-level gold matching. Retrieved CDEs were deduplicated by public identifier, retaining the highest-scoring version, so no public identifier can occupy more than one rank for a query. Ties in the BM25 score were broken deterministically by catalog order. Because BM25 has no learned parameters and no random initialization, the procedure is fully deterministic and was executed once; no seed average or standard deviation is reported for it.

For BM25 we swept the same canonical grid of four query representations by ten CDE text recipes used elsewhere in this work, evaluating every cell on Validation and freezing the best-performing cell before any evaluation set was touched. Selection ranked cells by Validation Recall@5 and then MRR@100. The selected configuration pairs the raw query text with its permissible-value summary (Q3) with the SN + LN + PQT + PV CDE recipe, which concatenates short name, long name, preferred question text and permissible-value summary in labelled form. Test and the distribution-shifted holdout sets played no part in this selection.

#### S5.2 CDE Match-Fuzzy keyword retrieval

CDE Match-Fuzzy searches CDE metadata with the exact, like and reverse-like rules of the documented matching flow, and adds character-n-gram and token-overlap fuzzy signals over DEC long name, long name, preferred question text. Each rule contributes candidates to a per-rule depth of 500, the fuzzy stage retains its top 1000 candidates, and the merged, deduplicated list is truncated to the top 10 candidates that enter the candidate pool. Evidence from several rules is aggregated per candidate, candidates are deduplicated by CDE public identifier, and exactly equal scores are broken deterministically by ascending public identifier.

Candidate depth. CDE Match-Fuzzy used a per-rule depth of 500 for the six evaluation datasets and 100 for Validation selection before truncation to the top 10 merged candidates. On Validation, the two settings produced identical top-10 lists for 99.6% of queries and no difference in gold inclusion at ranks 5, 10 or 20; all final feature evidence used depth 500.

Eligibility. Retrieval is restricted to the frozen production CDE catalog (62,976 CDEs; eligibility mode production_cde_match), which excludes retired, archived and non-production administrative records.

#### S5.3 Candidate-pool construction and selection of K

The reranker operates on a bounded candidate pool rather than the full catalog. For each query, candidates retrieved by FT-MPNet were combined with candidates from the keyword/fuzzy retriever, and the union was deduplicated by CDE public identifier. The nominal pool size K is the sum of the two retrieval depths. To choose K we held the keyword/fuzzy depth fixed at its deployed value of 10 and swept the bi-encoder depth, giving nominal pool sizes of K = 20, 30, 40, and 60. Because the two retrievers frequently return the same CDE, realized pools are often smaller than nominal K: at K = 30 the mean realized pool contains 27.6 candidates, and roughly 88-90% of queries yield fewer than 30 distinct candidates.

**Figure S4:**
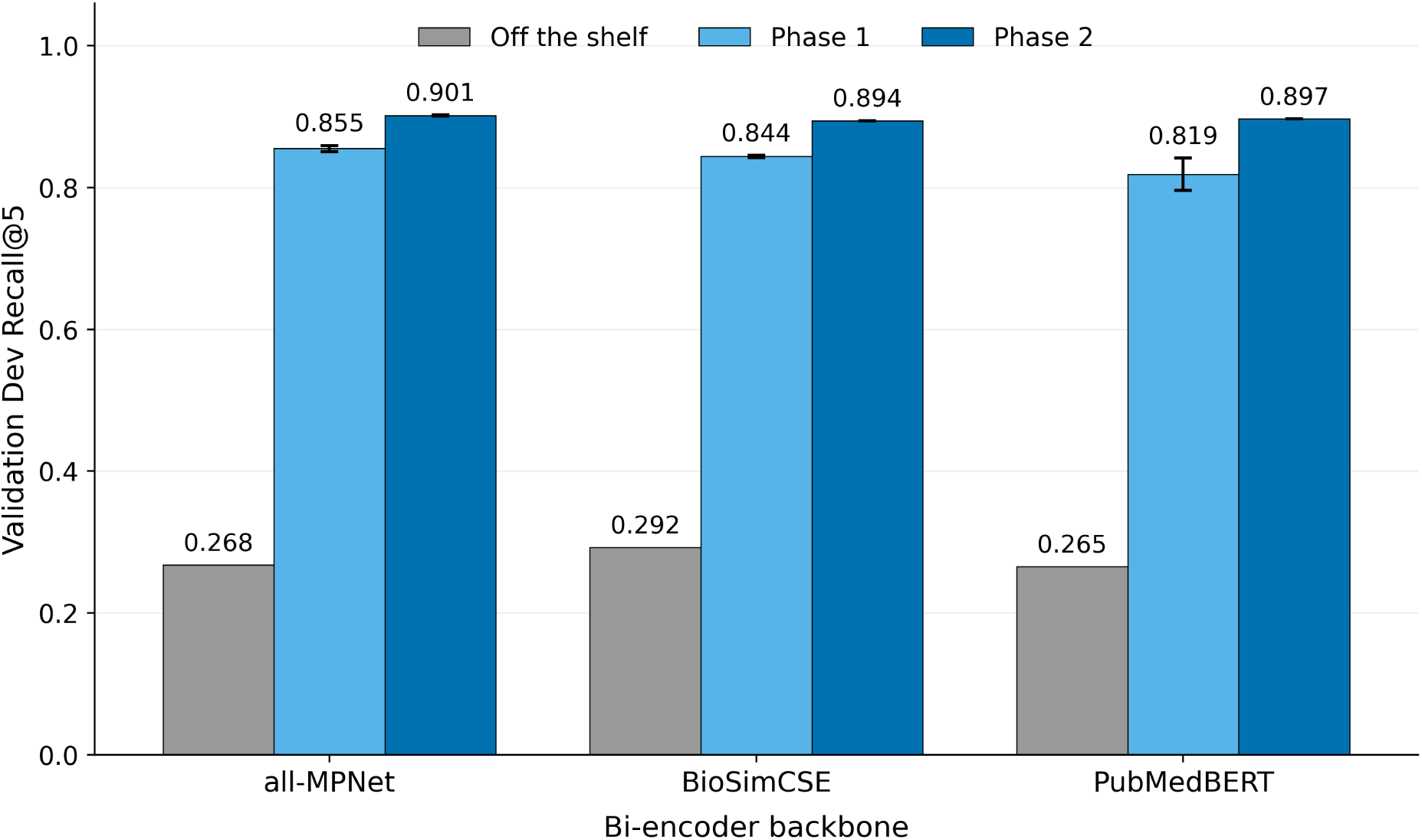
Validation Recall@5 for all-MPNet, BioSimCSE and PubMedBERT across off-the-shelf inference, Phase 1 fine-tuning and Phase 2 refinement. Fine-tuned stages show the two-seed mean with the sample standard deviation; off-the-shelf inference is deterministic and is shown without an error bar.

Pools were compared using ceiling Recall@5, defined as the proportion of queries for which at least one accepted gold CDE was present anywhere in the candidate pool. Because Recall@5 is hit-based, a gold CDE present anywhere in the pool could in principle be placed within the top five by an ideal reranker, so this quantity is equivalently the maximum Recall@5 achievable by an oracle reranker operating on that pool. It therefore bounds the performance of any reranker built on the pool and is independent of the reranker itself.

K was selected on Reranker Training; Validation was used only as descriptive confirmation and played no part in the choice. Ceiling Recall@5 rises from 0.9828 at K = 20 to 0.9888 at K = 30 on Reranker Training and then flattens, gaining only a further 0.0036 between K = 30 and K = 60 while roughly doubling the pool. We selected K = 30, at the onset of this plateau, as a coverage-versus-computation trade-off; Validation gives the same ceiling Recall@5 of 0.9888 at K = 30. This is a practical operating point rather than a statistically identified elbow, and no significance claim is made.

#### S5.4 Cross-encoder training and selection

Three cross-encoder backbones were compared: ncbi/MedCPT-Cross-Encoder, BAAI/bge-reranker-base, cross-encoder/ms-marco-MiniLM-L-6-v2. Training pairs were drawn from the corrected, production-eligible candidate pool, so no candidate carried empty text and no ineligible CDE was scored.

All backbones were trained under one frozen protocol (BCE pointwise, 2 epochs, learning rate 2e-5, batch size 32, maximum length 512, seed 20260527, trained on the full train split), with the positive class weighted by the ratio of negative to positive pairs, and were compared on Validation by Recall@5. FT-MedCPT was selected with Validation Recall@5 0.912, ahead of FT-BGE (0.904) and FT-MiniLM (0.874).

**Figure S5:**
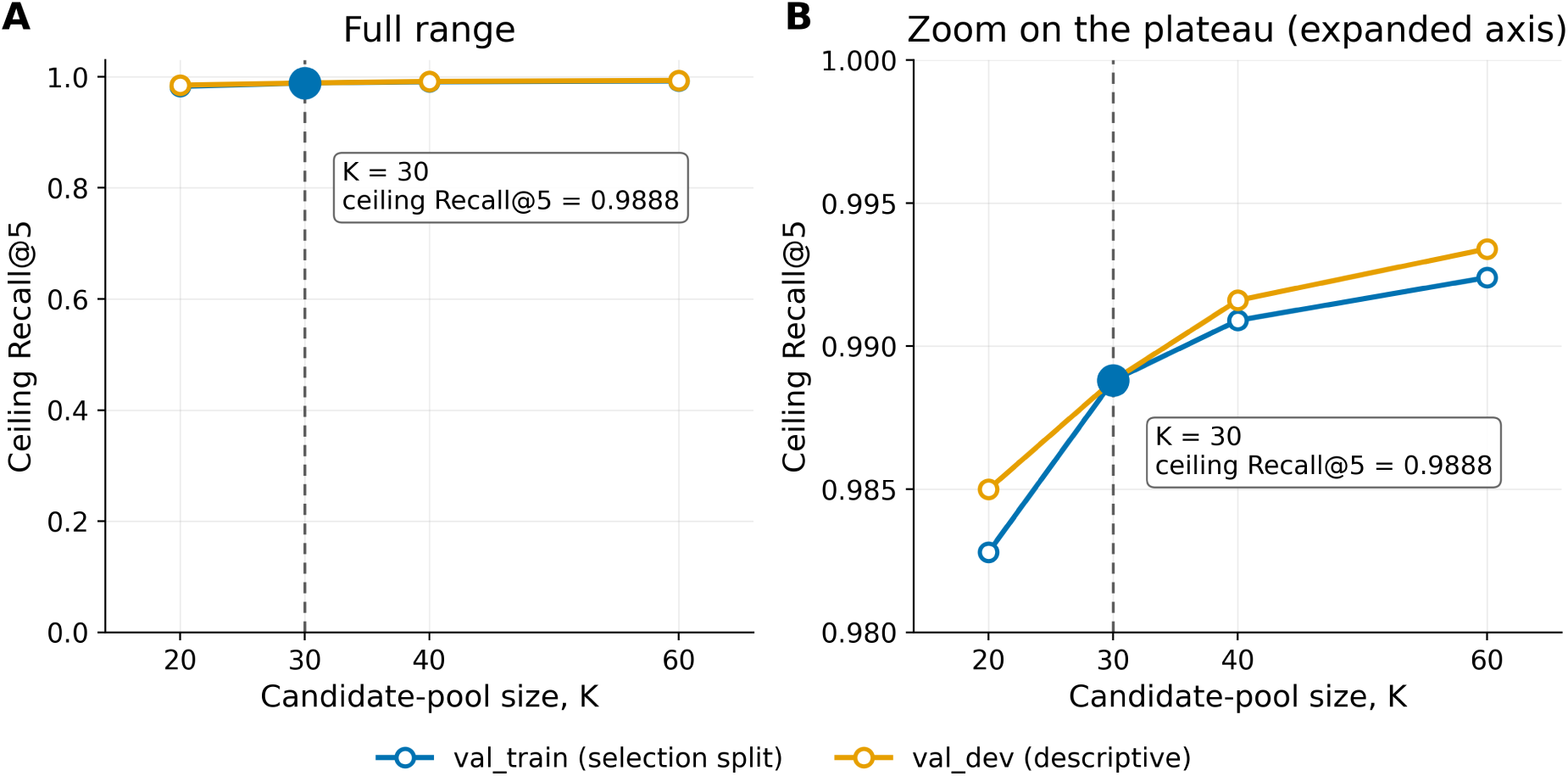
Ceiling Recall@5 as a function of nominal candidate-pool size K. Pools combine fine-tuned MPNet retrieval with a fixed keyword/fuzzy depth of 10. K was selected on Reranker Training; Validation is descriptive confirmation only. K = 30 was selected.

### S5.5 HGBC reranker

The final ranking model was a histogram-based gradient-boosting classifier (HGBC) implemented in scikit-learn. For each query, it scored the union of the top 20 FT-MPNet candidates and the top 10 CDE Match-Fuzzy candidates after deduplication by CDE public identifier. This produced a candidate pool of at most 30 CDEs per query, with a mean size of approximately 28. Candidate pools were constructed in the same way for training, validation, and evaluation, without adding the gold CDE when it was absent.

The HGBC was trained on the Reranker Training split, which contained 3,946 queries. Together, their candidate pools yielded 109,141 query–candidate pairs. A pair was labeled positive when the candidate’s public identifier matched any accepted gold CDE for that query; all other pairs were labeled negative. The training data contained 4,109 positive pairs, approximately 3.7% of the total. Queries with multiple accepted gold CDEs could contribute more than one positive pair. No negative sampling, class weighting, or sample weighting was used.

Each query–candidate pair was represented by 117 features in seven groups: retrieval-source indicators; FT-MPNet scores and ranks; keyword exact- and synonym-tier scores; permissible-value overlap; query and CDE text characteristics; keyword-rule evidence; and FT-MedCPT scores and ranks. Features that were not defined for a particular candidate—for example, a keyword score for a candidate retrieved only by FT-MPNet—were retained as missing values. The HGBC handles these values directly by learning, at each tree split, which branch missing values should follow. No global feature normalization or categorical features were used in the final model.

Hyperparameters were selected from 16 combinations of boosting iterations {200, 500}, maximum tree depth {3, 5}, learning rate {0.05, 0.1}, and minimum samples per leaf {10, 30}. Selection used Recall@5 on Validation, and no test or external evaluation dataset was used for training or model selection. The selected configuration used 200 boosting iterations, maximum depth 3, learning rate 0.05, minimum samples per leaf 30, and random seed 42. A prespecified follow-up comparison selected the feature set without query-provenance variables. Content-level training–evaluation overlap is characterized separately in Section S1.5 and Table S6.

At inference, candidates were ranked within each query by their predicted probability of being a gold CDE. Exact score ties were broken by ascending CDE public identifier.

### S5.6 Candidate-pool coverage and retrieval complementarity

**Table S5.**
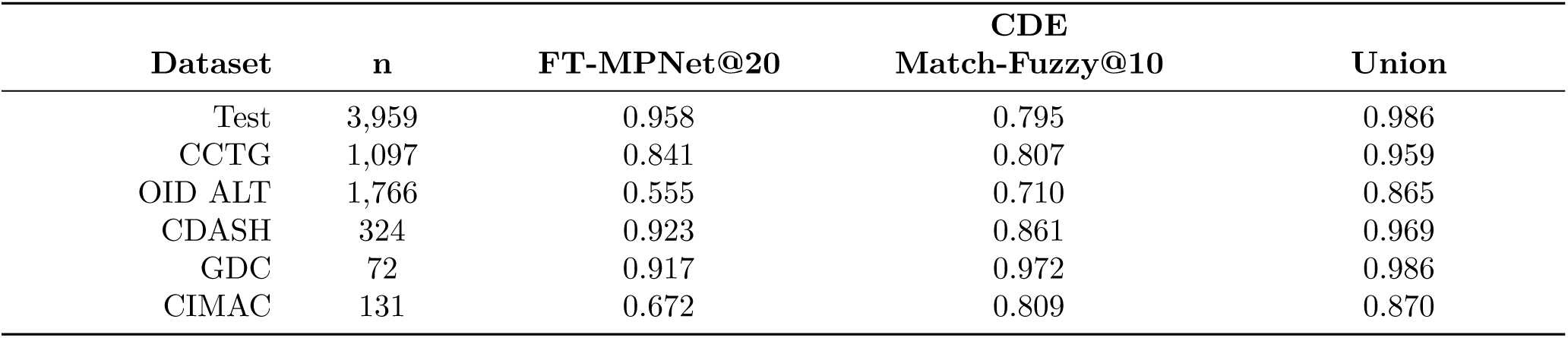
Candidate-pool gold coverage by retrieval arm. Coverage is the proportion of queries whose accepted gold CDE occurs anywhere in the stated candidate set, matched at the CDE public-identifier level; n is the denominator. The union is the fixed candidate pool used by the final reranker.

Coverage differed by retrieval method and dataset, but their union improved on the better individual retriever in every dataset, reaching 0.865–0.986.

### S5.7 Contribution of feature groups to final ranking

**Figure S6:**
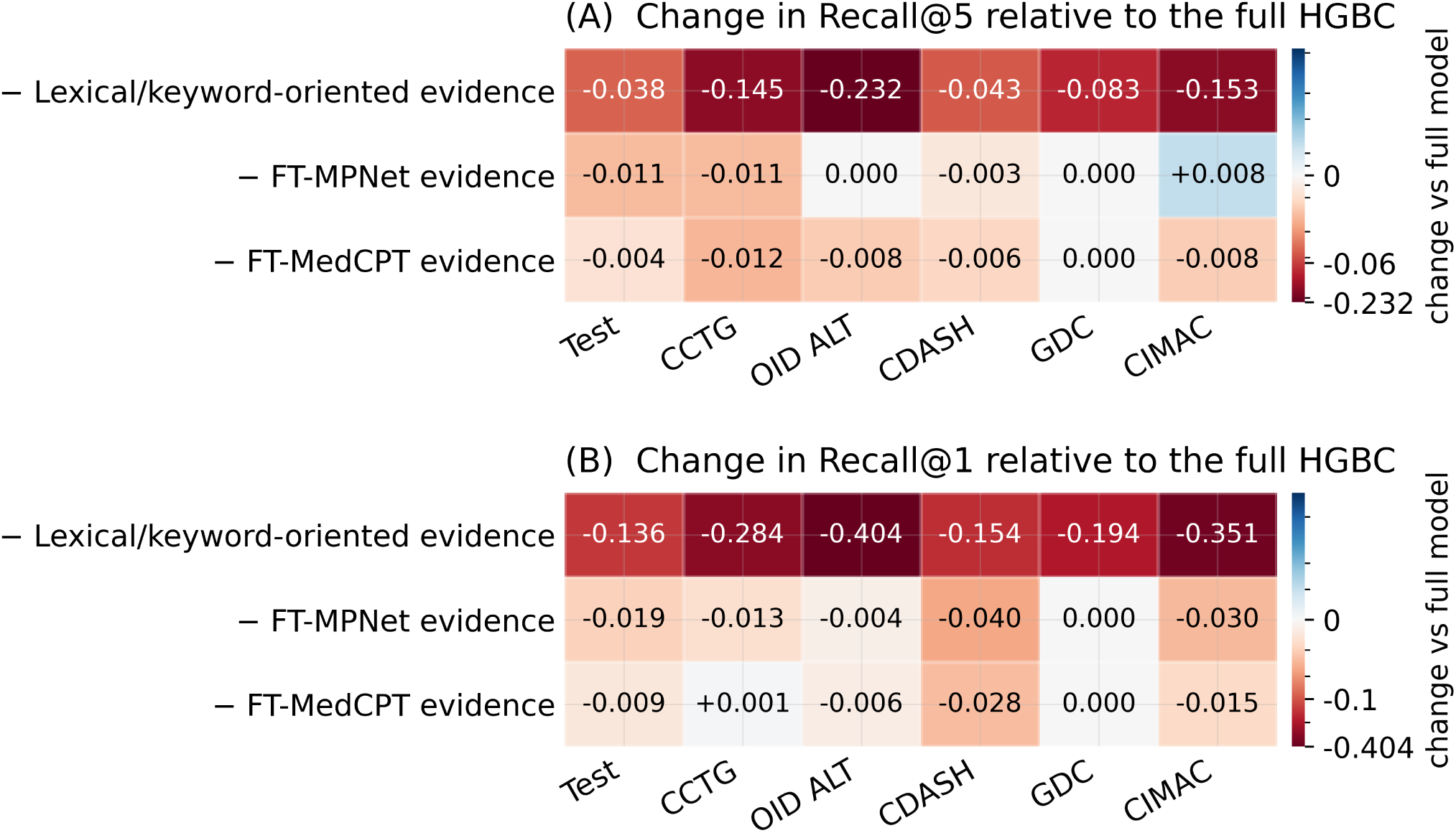
Broad evidence-family ablation of the final HGBC. Change in Recall@5 (A) and Recall@1 (B) after retraining the HGBC with one of three evidence families removed: lexical/keyword-oriented (98 features), FT-MPNet (13) and FT-MedCPT (6), which together partition all 117 features. All conditions use the identical fixed Stage-1 candidate pool, so the experiment measures contribution to ranking within the pool rather than to candidate generation. Cells give the change from the full 117-feature model; negative values (red) indicate lower recall. The colour scale is centred at zero and compressed beyond the largest neural-family effect so that the smaller effects remain visible; exact values are printed in every cell.

To isolate the evidence used for final ranking from candidate generation, we retrained the HGBC after removing each of three feature families while holding the Stage-1 candidate pool fixed. Removing lexical/keyword-oriented evidence produced the largest losses across datasets, whereas removing FT-MPNet or FT-MedCPT evidence produced smaller, dataset-dependent changes. This suggests that once a set of plausible candidates has been retrieved, detailed lexical cues such as identifiers, abbreviations, qualifiers, token overlap, and permissible-value agreement are especially useful for distinguishing the correct CDE. The lexical family contains multiple overlapping signals, so its contribution reflects their combined effect rather than any single keyword feature.

### S6. Sensitivity and robustness analyses S6.1 Training-evaluation overlap sensitivity

Table S6 evaluates whether removing repeated training–evaluation mappings changes the reported method performance.

**Table S6.** Sensitivity of Recall@5 to removing external-evaluation queries whose normalized text and gold CDE both recur in training. Filtered values are computed by rescoring the same saved rankings against the leakage-filtered query lists. The internal Test split is in-distribution by design and is not included.

| Dataset | Method | n (all) | Recall@5<br>(all) | n (filtered) | Recall@5<br>(filtered) | $\Delta$ |
| --- | --- | --- | --- | --- | --- | --- |
| CCTG | Final reranker | 1,097 | 0.909 | 1,089 | 0.909 | +0.0002 |
| CCTG | Python approximation to NCI CDE Match | 1,097 | 0.739 | 1,089 | 0.738 | -0.0010 |
| CCTG | BM25 | 1,097 | 0.392 | 1,089 | 0.394 | +0.0020 |
| OID ALT | Final reranker | 1,766 | 0.832 | 1,759 | 0.832 | -0.0007 |
| OID ALT | Python approximation to NCI CDE Match | 1,766 | 0.770 | 1,759 | 0.769 | -0.0009 |
| OID ALT | BM25 | 1,766 | 0.176 | 1,759 | 0.176 | +0.0001 |
| CDASH | Final reranker | 324 | 0.920 | 310 | 0.919 | -0.0004 |
| CDASH | Python approximation to NCI CDE Match | 324 | 0.750 | 310 | 0.748 | -0.0016 |
| CDASH | BM25 | 324 | 0.710 | 310 | 0.729 | +0.0192 |
| GDC | Final reranker | 72 | 0.972 | 68 | 0.971 | -0.0016 |
| GDC | Python approximation to NCI CDE Match | 72 | 0.986 | 68 | 0.985 | -0.0008 |
| GDC | BM25 | 72 | 0.542 | 68 | 0.515 | -0.0270 |
| CIMAC | Final reranker | 131 | 0.802 | 113 | 0.779 | -0.0228 |
| CIMAC | Python approximation to NCI CDE Match | 131 | 0.679 | 113 | 0.637 | -0.0422 |
| CIMAC | BM25 | 131 | 0.160 | 113 | 0.150 | -0.0099 |

### S6.2 Exact-match allowance sensitivity

Both lexical methods used the same deterministic query-level mask to limit direct exact lookup of the gold CDE. The exact-match rules search the CDE long name, question-text strings in its reference documents—including preferred and alternate question text—and alternate names. A query harvested from one gold-linked metadata field can be identical to a string stored in another, so the mask covers all exact-match-eligible fields rather than only the field from which the query originated.

Queries were assigned to nested allowance subsets using a SHA-1 hash of the query identifier with seed 42. For a masked query, only strings belonging to the gold CDE that were identical to the normalized query were withheld. The query itself, other catalog entries, the gold CDE’s nonidentical metadata, non-exact matching rules, and permissible values were unchanged. The Python approximation to NCI CDE Match and CDE Match-Fuzzy therefore received the same exact-match availability at every allowance rate.

The 70% operating rate was motivated by CIMAC, where 92 of 131 queries whose gold CDE was present in the catalog had an exact match when all eligible metadata were available (70.2%). We therefore used a 70% allowance for the four caDSR-derived evaluation sets and a 100% allowance for GDC and CIMAC, which were assembled independently and do not create the same circularity.

The fixed HGBC trained at the 70% allowance was substantially less sensitive to allowance changes than either lexical method. Across the full 0-100% range its Recall@5 varied by 0.075 on Test and 0.164 on CDASH, whereas the two lexical methods varied by 0.40-0.96 on every dataset. At the 0% allowance, where every exact-matching string of the gold CDE was withheld, the fixed reranker still reached 0.920 on Test and 0.818 on CDASH, compared with 0.151-0.556 for the lexical methods. Its performance also remained close to that of HGBC models retrained separately at each allowance, differing by at most 0.02 Recall@5 in 20 of 24 dataset-rate conditions; the largest difference was on CDASH at the 0% allowance, where retraining recovered 0.065 Recall@5. Retraining therefore provided limited additional benefit under most conditions rather than none. OID ALT remained the most allowance-sensitive dataset, with Recall@5 falling from 0.832 at the 70% allowance to 0.463 at 0%. Much of that loss was a limit of candidate generation rather than of ranking: the proportion of OID ALT queries whose accepted gold CDE entered the candidate pool at all fell from 0.865 at the 70% allowance to 0.571 at 0%, where 758 of 1,766 gold CDEs were absent from the pool. No reranker can recover a gold CDE that is not in the pool. At the 100% allowance all methods converged near ceiling, so that condition is an upper anchor rather than a discriminating comparison.

**Table S7.**
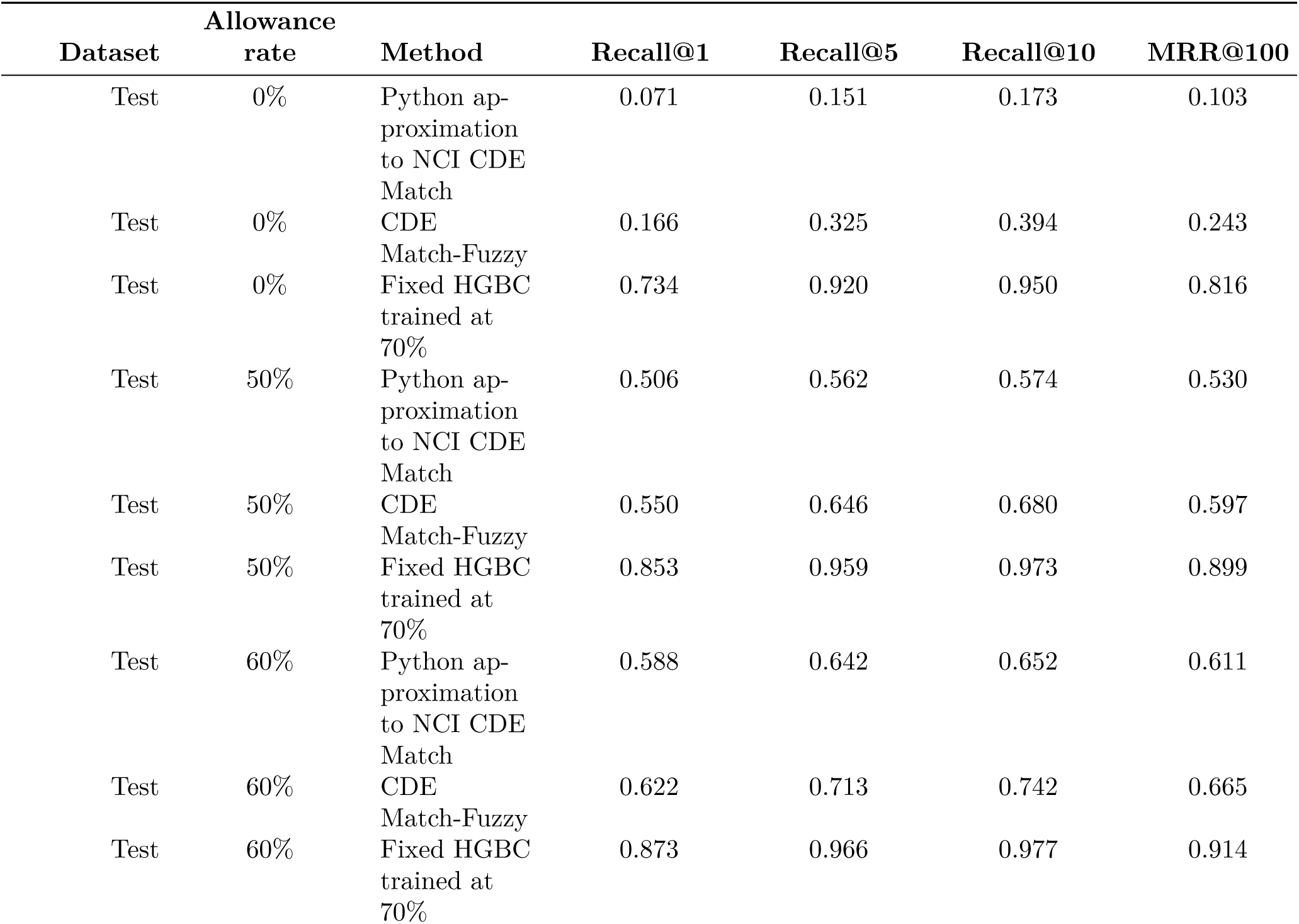
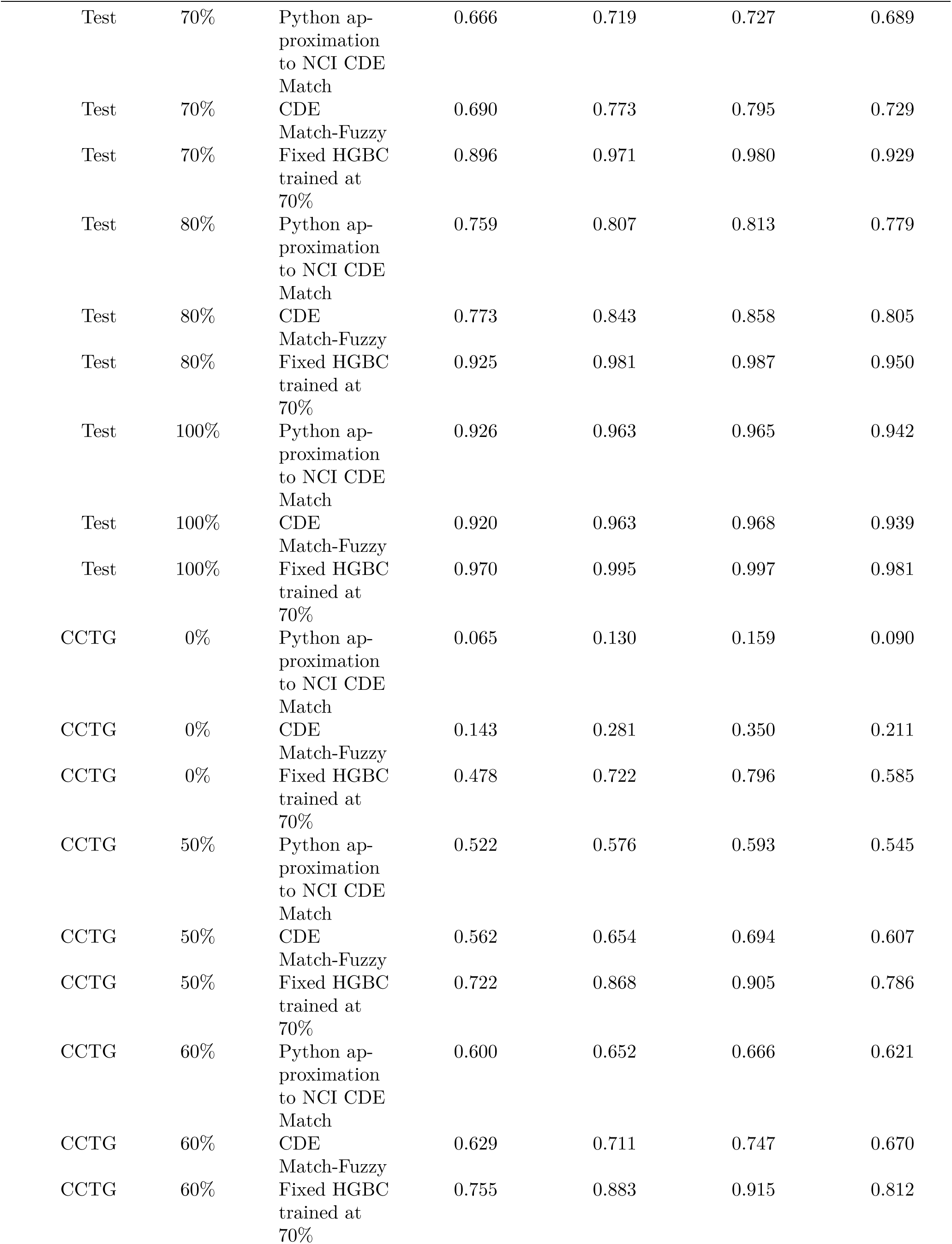
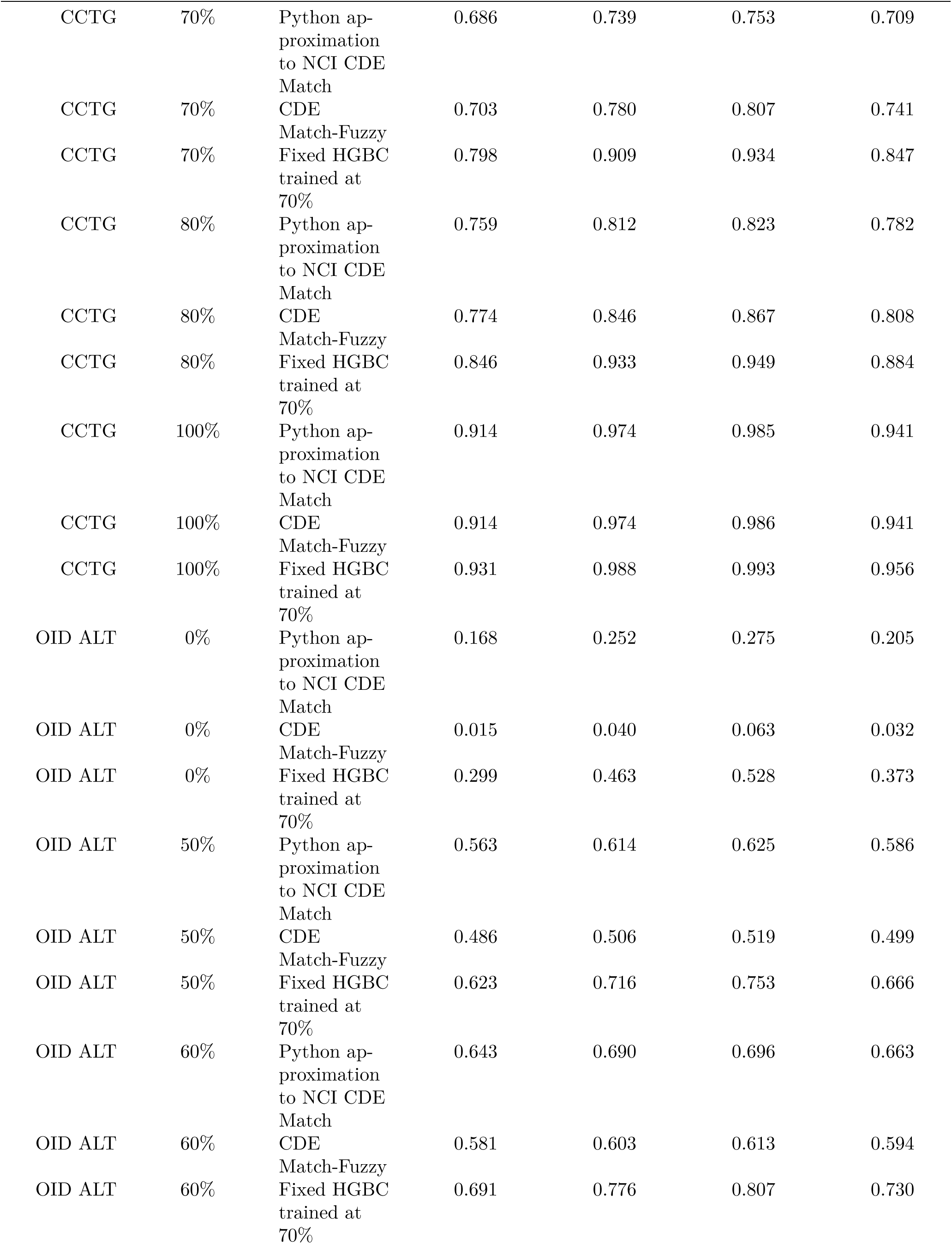
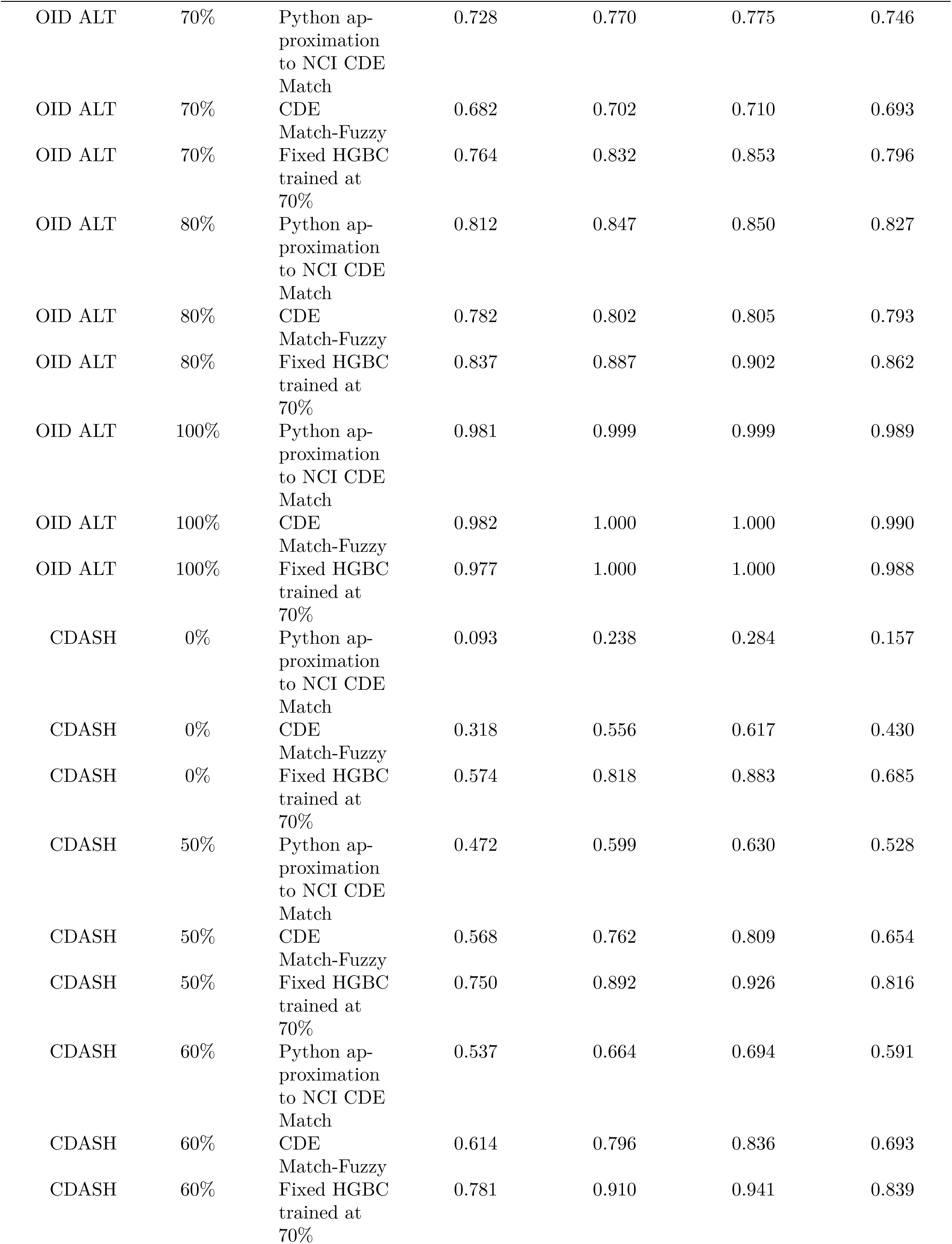
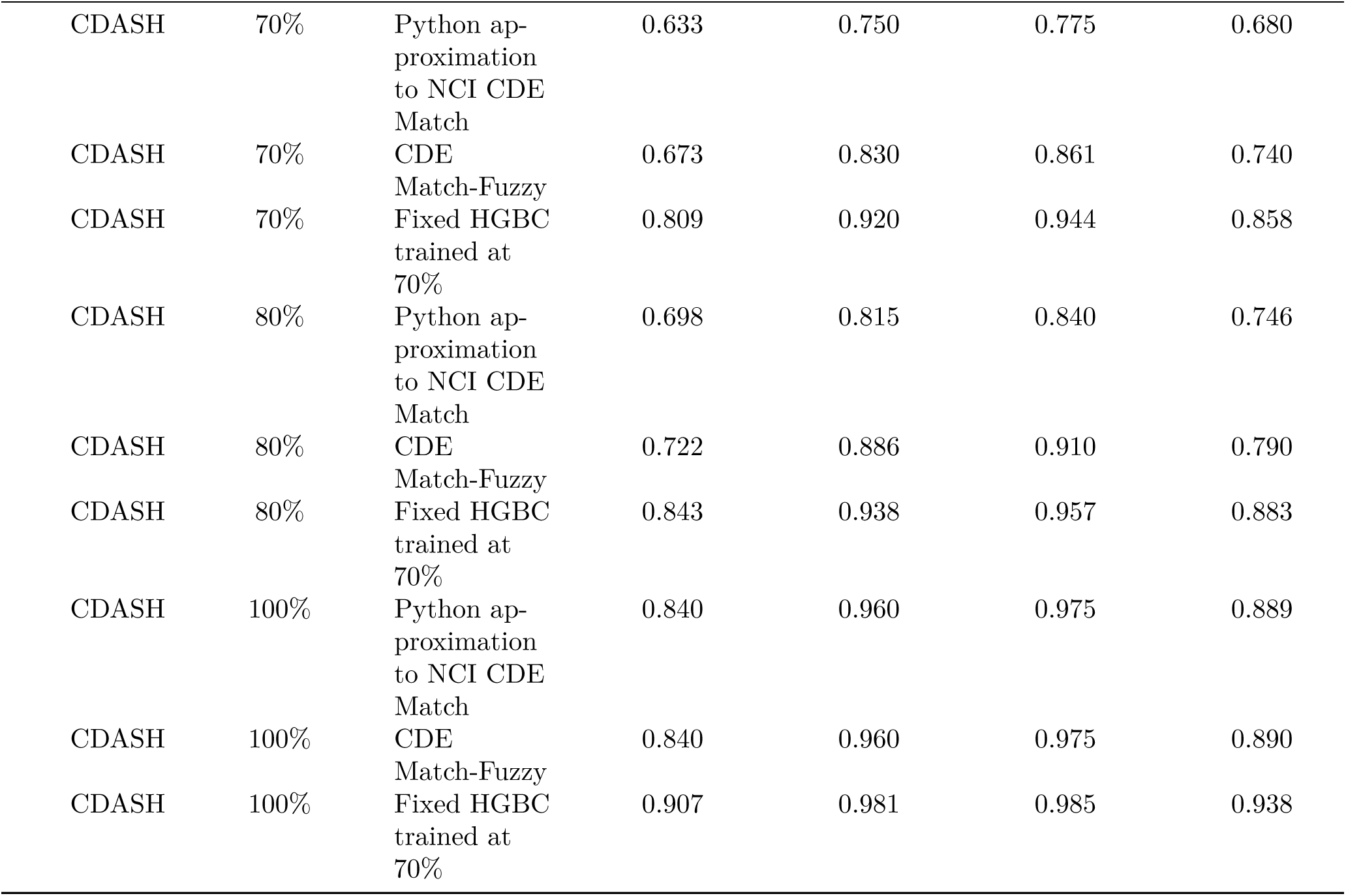
Exact-match allowance sensitivity for the Python approximation to NCI CDE Match, CDE Match-Fuzzy, and the fixed HGBC trained at 70% allowance: Recall@1, Recall@5, Recall@10, and MRR@100.

**Figure S7:**
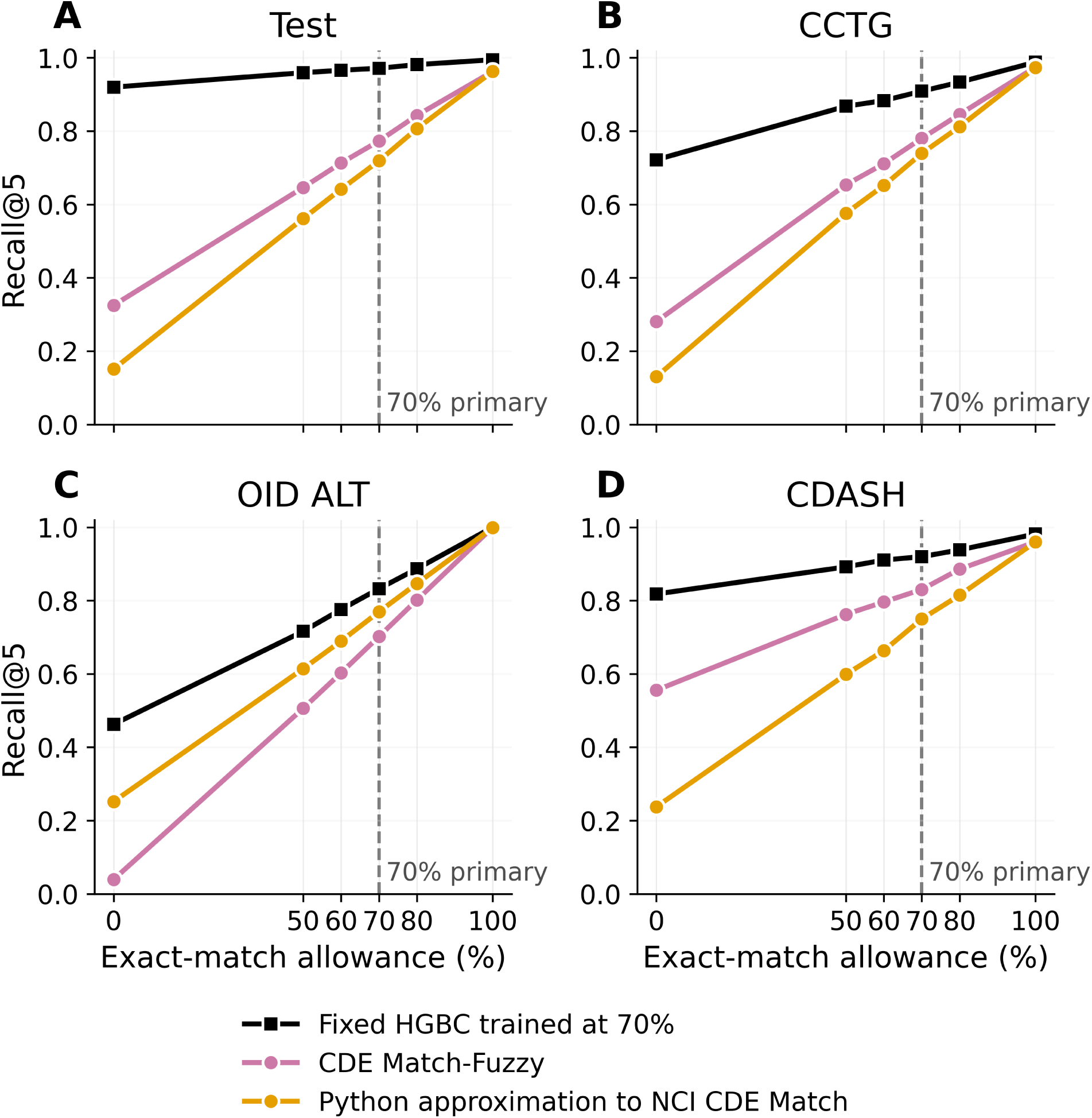
Recall@5 as a function of exact-match allowance on the four caDSR-derived evaluation sets. The HGBC curve uses the unchanged reranker trained at the 70% allowance. At each inference allowance, CDE Match-Fuzzy candidates, merged candidate pools, and allowance-dependent features were regenerated, while FT-MPNet, FT-MedCPT, and HGBC model weights remained fixed. The 70% operating point used in the primary analysis is marked.

**Figure S8:**
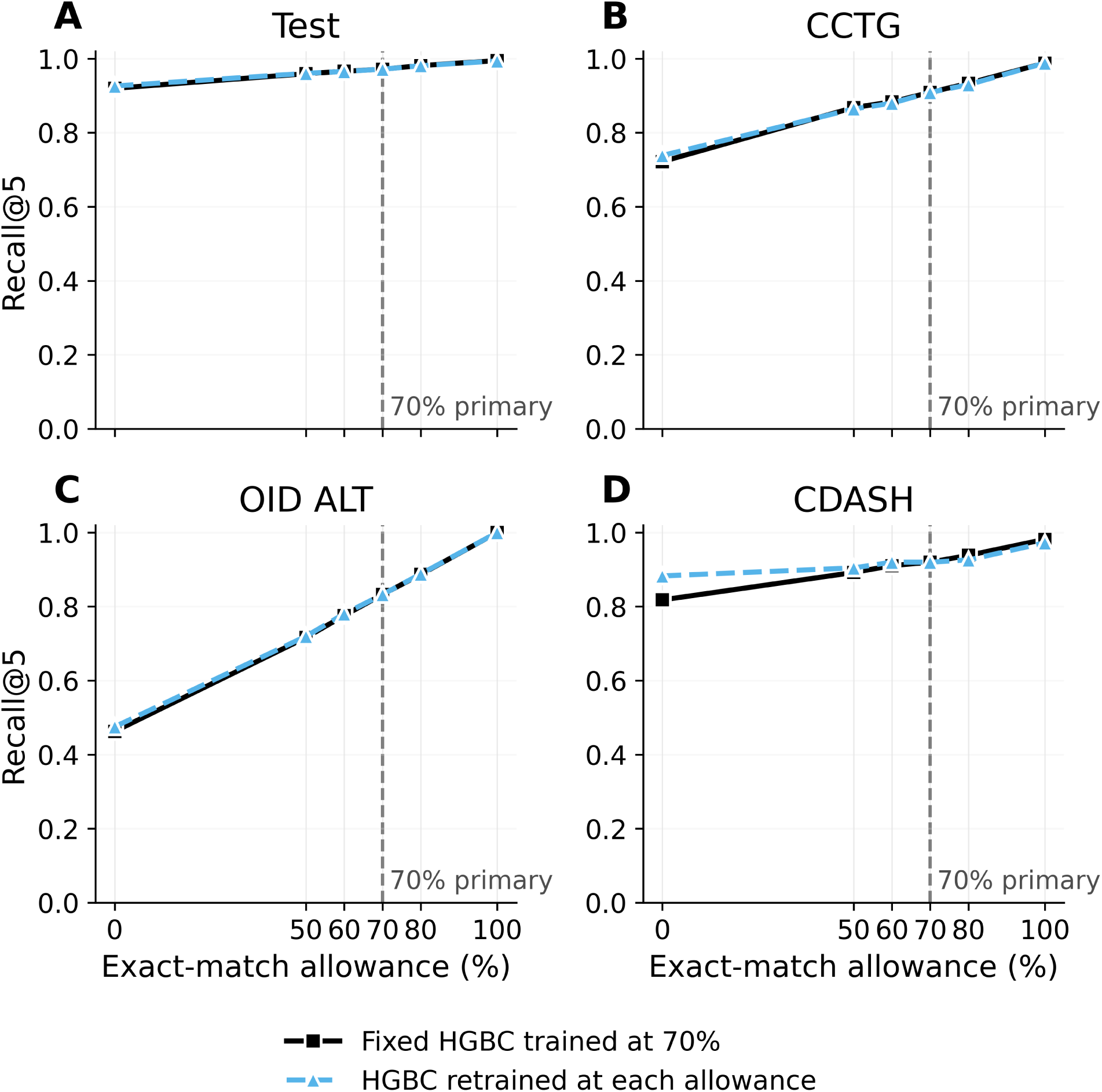
Recall@5 for the fixed HGBC trained at the 70% allowance and HGBC models retrained separately at each allowance. Candidate pools and allowance-dependent features were generated at the indicated allowance for both analyses; FT-MPNet and FT-MedCPT remained fixed. The fixed-model analysis represents inference-time mismatch in a deployed pipeline, whereas the retrained analysis measures adaptation of the HGBC training procedure.

